# Injury Induced Connexin 43 Expression Regulates Endothelial Wound Healing

**DOI:** 10.1101/2025.02.24.639946

**Authors:** Meghan W. Sedovy, Mark C. Renton, Kailynn Roberts, Xinyan Leng, Clare L. Dennison, Melissa R. Leaf, Paul D. Lampe, Angela K. Best, Brant E. Isakson, Scott R. Johnstone

## Abstract

Endothelial cell (EC) injury is a major contributing factor to vascular surgical failure. As such, understanding the mechanisms of endothelial healing is essential to the development of vascular therapeutics and procedures. Gap junctions formed by connexin 43 (Cx43) are implicated in regulating skin wound healing, but their role in endothelial healing is unknown. Secondary analysis of RNAseq data from *in vivo* injured mouse aortas (GEO: GSE115618), identified significant Cx43 upregulation in EC post-injury. We developed a novel in vivo model of EC injury using mouse carotid artery ligation to test the role of Cx43. We identified that EC immediately adjacent to the wound edge upregulate Cx43 protein expression, predominantly at cell-cell junctions. We show significantly delayed EC healing in a mouse model of inducible EC-specific Cx43 deletion (EC-Cx43 KO) at 24 hr post ligation. Single cell RNAseq analysis of 10,829 cells from 18 hr injured EC-WT and EC-Cx43 KO carotids revealed a Cx43-associated reduction in enrichment of EC pathways associated with migration, proliferation, and ERK/MAPK signaling pathways. Finally, the importance of Cx43 phosphorylation on EC healing was tested in mice with single-point alanine mutations (phospho-null) in known phosphorylation sites that alter Cx43 channel assembly and opening. Mice containing alanine mutations at ERK phosphorylated Cx43 serines (Cx43^S255/262/279/282A^) reduces healing rates similar to EC-Cx43 KO. These data suggest that EC injury-induced Cx43 upregulation, and subsequent Cx43 gap junction-mediated cell-to-cell communication are required for normal EC migration during wound healing after vascular injury.

**New and Noteworthy:** These findings demonstrate for the first time that mechanical injury to large artery endothelium induces the expression of gap junction protein Cx43. This upregulation improves migratory and proliferative capacity of endothelial cells at the wound edge, facilitating timely wound closure. This phenomenon is dependent on appropriate gap junction function and turnover.

## Introduction

Cardiovascular disease is the leading cause of death and disease globally (1). One of the most prevalent forms of cardiovascular disease, coronary artery disease, affects approximately 20.5 million Americans (2). Consequently, procedures designed to correct these coronary blockages, such as percutaneous coronary interventions (PCI) and coronary artery bypass grafts, are among the most frequently performed cardiovascular surgeries (2). More than 160,000 PCI and 434,000 coronary artery bypass grafts are conducted annually in the US (2). Despite their widespread use, these surgeries often fail in the first few years following surgery (3, 4), and vascular endothelial cell (EC) damage is thought to be a key contributor to early failure (5, 6).

The innermost lining of blood vessels consists of a single layer of ECs. In large conduit arteries, such as the coronary artery, ECs play a crucial role in protecting the vessel from excessive platelet aggregation, inflammation, and disease (7). During vascular surgery, mechanical damage to ECs leads to cell death and denudation of EC from large portions of the vessel wall (8, 9). As a result, vessels are at increased risk of thrombosis, neointima formation, and neo-atherosclerosis (10–13). To date, various techniques have been employed to preserve EC integrity during vascular procedures or to promote reendothelialization after damage occurs. We recently reported that saphenous vein storage during coronary artery bypass surgery can significantly impact EC coverage. Our study and others show that standard heparinized saline causes EC damage and increases oxidative stress, whereas specially formulated solutions and autologous patient blood improve EC health and retention (9, 14–16). Despite this, the adoption of these surgical techniques is not widespread, and a degree of endothelial damage and loss was shown to be inevitable regardless of the preparation performed (9). Therefore, understanding mechanisms controlling the timely healing of the vascular endothelial layer is critical for surgical success and informing future therapies.

Efforts to enhance reendothelialization have included EC-capture stents designed to attract endothelial progenitor cells from the bloodstream to the denuded stent site, yet clinical trial evidence has not demonstrated superior efficacy of these stents compared to the more commonly used drug-eluting stent (17, 18). This may be because studies demonstrate that wound healing relies on the proliferation and migration of wound-adjacent EC, rather than recruitment of EC progenitors (19, 20). In large arteries, vascular ECs are typically quiescent, exhibiting minimal proliferation and cell turnover (21–23). Following vascular injury, wound-adjacent EC become activated, increasing transcription factors linked with proliferation (e.g. Myc/FoxM1/FoxO1) and regulators of cell-to-cell junction integrity and signaling (FosL2 and Hippo signaling) (20). Many junctional complexes are retained during large artery EC healing and it is possible that these proteins are needed to regulate cell proliferation, migration, or both (20, 24, 25).

Gap junction channels connect adjacent cells and allow cell-to-cell exchange of ions and small metabolites (<1.2 kDa) to coordinate tissue functions. Gap junctions formed by connexin (Cx) proteins are widely expressed, although only Cx37, Cx40, Cx43, and Cx45, are reportedly expressed in vascular cells (26–32). In large artery EC, Cx37 and Cx40 are the most widely expressed, playing roles in signal transduction along the vessel wall. While Cx43 is expressed in EC of resistance arteries (33), in larger conduit arteries, Cx43 expression is limited to vascular branch points and regions of turbulent flow with elevated mechanical stress (27, 28, 34–36). Cx43, is the most extensively studied connexin and has been widely associated with the control of cellular proliferation and migration control. However, studies have reported both increases and decreases in cell proliferation following Cx43 knockdown or alteration and the exact mechanisms remain to be fully elucidated (37–41). Studying Cx43 regulation in large artery EC *in vitro* does not accurately replicate the *in vivo* setting, as Cx43 expression is highly upregulated in culture conditions (42). Despite this, *in vitro* studies have demonstrated that knockdown of Cx43 in cultured EC reduces EC proliferation and migration (42, 43). These data in conjunction with data from skin wounding models, suggest a likely role for Cx43 gap junction channel signaling in EC healing.

Gap junction channel, assembly, trafficking and opening are highly regulated by phosphorylation of the connexin c-terminus (44, 45). The Cx43 c-terminus contains 29 serines, threonines and tyrosines that are differentially phosphorylated by multiple kinases (e.g. PKC, MAPK, and AKT) which dynamically control channel gating and gap junction intracellular communication (46–56). Specifically, phosphorylation of connexin c-terminus amino acids governs the channel’s aggregation into larger gap junction plaques, its removal from the membrane, and its interactions with other intracellular proteins (45, 57–59). Cx43 expression, gap junction activity, and protein-protein interactions modulate cell migration and proliferation during normal development and pathological conditions (38, 60–71). However, the specific role of Cx43 appears to vary depending on cell type, condition, or disease (37, 38, 60–65, 72–74). Given that EC expresses Cx43 during periods of stress and Cx43 modulates cell proliferation and migration, it is likely that Cx43 and its phosphorylation regulated functions are involved in EC wound healing. However, the relationship between endothelial injury and Cx43 *in vivo* has not been thoroughly explored.

In the present study, we identified a significant induction of EC Cx43 expression and gap junction formation *in vivo* following vascular injury. Single-cell RNA sequencing in Cx43 knockout EC revealed that injury-induced Cx43 expression promotes EC wound healing through the expression of genes that facilitate cell migration/proliferation. Finally, we show that Cx43 control of EC wound healing functions is dependent on gap junction regulating Cx43^S255/262/279/282^ phosphorylation.

## Materials and Methods

### Animals

All animal protocols and care procedures were approved by the Virginia Tech Animal Care and Use Committee. Both male and female mice were included in all studies, with sex assessed as a biological variable. C57BL6/J mice (JAX strain #000664) and CDH5^(PAC)CreERT2^ mice (75) and Cx43^fl/fl^ mice (JAX strain #008039) (76) on a C57BL6/J background were purchased from Jackson Laboratory and bred in house. Heterozygous CDH5^(PAC)CreERT2^ mice were bred with Cx43^fl/fl^ mice (JAX strain #008039) to produce CDH5^(PAC)CreERT2^Cx43^fl/fl^ (EC-Cx43 KO) and littermate and Cre negative Cx43^fl/fl^ mice and (EC-WT) mice. Additionally, mice expressing phospho-null amino acid mutations (serine to alanine) on the c-terminal tail of Cx43 that prevent phosphorylation were used to test the impact of Cx43 gap junctions. Phosphorylation at sites associated with protein kinase C (PKC) (Cx43^S368A^) (77), mitogen-activated protein kinase (MAPK) (Cx43^S255/262/279/282A^) (45, 78), and protein kinase A and B and (PKA/AKT) (Cx43^S365/373A^) respectively were used. All studies were performed in adult mice aged 12-22 weeks.

### Tamoxifen Feeding

Induction of Cre was performed using tamoxifen feed (Inotiv 500 mg/kg tamoxifen) to avoid excess inflammation associated with injection (79). To ensure efficient knockdown in CDH5 strains, mice were fed tamoxifen diet ad libitum for four weeks followed by a two-week washout period during which the mice were fed regular chow (80). Tamoxifen diet consumption was monitored by tracking the weight of the mice, with wet food provided for the first week of feeding. Gene and protein knockout of Cx43 in EC was confirmed by PCR from tamoxifen-fed mouse aortas and by immunofluorescence in tamoxifen-fed injured mouse carotid EC, respectively (Fig 2B and C, Supplemental Fig. 4 and 6).

### Genotyping/PCR

DNA extracted from mouse tissue was used to confirm mouse genotypes using primers detecting CreERT2 (Generic Cre), CDH5 Cre, Cx43 flox, and/or GAPDH control. Probes published by Liao et al. (76) were used to detect Cx43 exon 2 deletion in CDH5^(PAC)CreERT2^, Cx43^fl/fl^ mice using DNA extracted from EC containing aortic tissue of tamoxifen fed mice, a 686 band bp in these experiments indicates Cx43 knockout. A list of primers used can be found in **Supplemental Table 1**.

### Temporary Carotid Ligation Endothelial Injury Surgery

Mice undergoing carotid endothelial injury surgery were anesthetized with isoflurane. The surgical site was cleared of hair and sterilized. Each mouse received 0.5 mL of extended-release analgesic (Ethiqa) before surgery. During the procedure, mice were hydrated with a subcutaneous injection of saline, kept warm on a heating pad, and lubricant applied to their eyes. A midline incision was made in the neck, followed by clearing of tissue and fat to expose the left carotid artery. A section of the carotid was fully isolated from surrounding tissue, and a 5-0 suture was placed underneath. The suture was secured, stopping blood flow for one minute before being untied, allowing blood flow to be restored. The incision was then closed, and the mouse was placed in a clean cage on a heating pad for recovery. At the end of the indicated time points, mice were euthanized by CO2 inhalation and cervical dislocation for harvesting of injured carotid artery tissue for specific experiments described below.

### Immunofluorescence

Carotid artery tissue collection for immunofluorescent staining was conducted by cardiac perfusion in euthanized mice with ∼3 mL heparin-saline solution, followed by ∼3 mL 4% paraformaldehyde (PFA) at a rate of 1 mL/min. Carotids for longitudinal sectioning were fixed in 4% PFA overnight, and embedded longitudinally for sectioning. Paraffin-embedded sections were deparaffinized with standard xylene procedures, and antigen retrieval was performed using a pressure antigen retrieval machine (Aptum Biologics retriever 2100) with universal epitope antigen retrieval buffer (Electro Microscopy Sciences #62719-20). Samples were blocked under antibody-specific conditions, followed by the addition of primary antibodies in a blocking buffer overnight, washing, then detected using fluorescent tagged secondary antibodies. FadeStop fluorescent mounting medium with DAPI (LifeSct #272L) was added prior to imaging. For en-face imaging, carotid arteries were cut open longitudinally before removal, then fixed in 4% PFA for 1 hr before proceeding with blocking, overnight staining with primary antibody, and 1 hr incubation with secondary antibody directly in 1.7 mL tubes. A detailed list of blocking conditions, primary and secondary antibodies, and concentrations can be found in **Supplemental Table 1.**

## Measurements of Wound Closure

All imaging was performed on a Nikon NIE confocal, with resonance scanning and FTP-2000 motorized X/Y/Z platform allowing for tiled and Z-stack images. Wound length was measured using NIS-Elements AR (version 5.42.04). For all wound healing experiments, EC were detected using immunofluorescent staining of the known EC marker vascular endothelial cadherin (VE-cad). EC wound area was considered the length in microns of vessel missing EC, demonstrated by a lack of VE-cad signal. Cx43 punctuates were measured using NIS-elements thresholding to identify individual puncta. The ratio of punctates to tissue area was calculated (punctates/area) and multiplied by 100.

### Western Blotting in Flushed Carotid Endothelial Cells (ECs)

Carotid EC were collected by perfusing euthanized mice with ∼3 mL heparin-saline, followed by a ∼3 mL saline flush to remove residual heparin at a rate of 1 mL/min. Carotids were isolated, and an incision made at the aortic junction with the carotid artery. A 25-gauge needle attached to a syringe containing 100 µL of NP-40 substitute (0.1%) protein lysis buffer with added protease and phosphatase inhibitors (**Supplemental Table 1**) was inserted through the incision into the carotid artery. A hemostat secured the needle, and the carotid artery was excised by cutting tissue around the clamped needle and at the distal end of the carotid. The free end of the carotid was placed over a 1.7 mL tube, and a small amount of lysis buffer was flushed through the artery. Lysis buffer was left in the carotid for one minute before the remainder was pushed through under pressure into the collection tube. Samples were immediately placed on ice, vortexed, and centrifuged at 14,000 g for 5 min at 4°C. The supernatant was transferred to a clean tube and stored at −20°C. The remaining flushed vessel (denuded vessel) was unclamped, lysed, and stored at −20°C. Western blotting of samples was performed using Thermo Fisher 10% Bis-Tris gels. Sample protein loading was confirmed with LI-COR Revert 700 total protein assay. VE-cad was used as a marker for EC, and transgelin (TAGLN) was used to detect the presence (or absence) of smooth muscle cells (contamination in EC only preps). Expression of Cx43 in EC samples was normalized to VE-cad expression.

### Carotid Collection and EC-enriched Digestion for Single Cell RNA Sequencing (scRNAseq)

Both EC-Cx43 KO and EC-WT mice were euthanized by CO2 overdose and cervical dislocation 18 hr post carotid EC injury and the injured carotid was exposed without perfusion to minimize time and disruption to the endothelial layer. The carotid was then removed from the mouse and dipped in a solution of heparin saline followed by a saline-only solution. Before digestion, carotids were stored in ice-cold EC media (Vascular Cell Basal Medium; ATCC #PCS-100-030) supplemented with all components of an Endothelial Cell Growth Kit (ATCC #PCS-100-041) excluding fetal bovine serum (FBS) while subsequent carotids were removed. To enrich for EC while collecting enough total cells for processing and sequencing, carotids were cut open lengthwise prior to removal to expose the EC layer for digestion. Carotids were kept whole and a short 20 min digest (described below) was performed.

Up to two carotids were digested in a 1 mL digestion solution of 0.2 mg/ml liberase TM (Roche #5401119001) and 60 U/ml DNase1 in supplemented EC media. Carotids were incubated at 37°C and 400 rpm in a benchtop orbital shaking heat block for 20 min, pipetting up and down with a 1 mL pipette every 5 min. Digestion was stopped with the addition of 1 mL stopping solution (2% FBS, 1mM EDTA, in supplemented EC media), after which samples of the same condition were combined (up to 5 at a time to reduce time to digestion). A 100 μm cell strainer was primed by flushing through 2 mL of supplemented EC media with 2% FBS. Cell solutions were filtered through the strainer and rinsed with 3 mL of supplemented EC media with 2% FBS. The flow through was centrifuged at 1000 rpm, 4°C for 10 min, followed by removal of the supernatant and resuspension of the cell pellet in 2 mL RBC lysis buffer (Invitrogen 00-4333-57) at room temperature for 5 min. RBC lysis was stopped using 10 ml supplemented EC media with 2% FBS, samples were centrifuged at 1000 rpm, 4°C for 10 min and the cell pellet was resuspended in 10% DMSO, 10% FBS in cold supplemented EC media. Single cell suspensions were slowly frozen at −80°C overnight in a cell freezer container, then transferred to liquid nitrogen until sequencing. A total of 15 carotids for EC-WT and 20 carotids for EC-Cx43 KO were pooled for use for scRNAseq by Novogene.

### Novogene scRNAseq Methods

Cell viability analysis, RNA integrity analysis, library preparation, sequencing, raw read count generation, and reference genome alignment were performed commercially by Novogene (Sacremento, CA, USA) using their Plant and Animal 10X Single Cell Transcriptome Sequencing pipeline. Full parameters, protocols, and quality control reports are available on request, but are explained briefly below:

### Sample and Library Preparation

Cryopreserved cell samples were thawed and processed by Novogene. Cell count and viability were assessed using a TC20 Automated Cell Counter (Bio-Rad, USA) with both samples achieving approximately 60,000 cells with an overall cell viability of >75%. Libraries were then constructed using the 10x Chromium Single Cell 3’ Kit with a target of 10,000 cells.

### Sequencing and Data Preprocessing

Sequencing was performed on the NovaSeq X Plus Series (Illumina, USA) platform with 150 bp paired end reads with a minimum of 150 G raw data per sample. Raw sequencing data in FASTQ format were pre-processed with Cell Ranger (v7.0.0) using default parameters, and subsequent reads were aligned to the mouse genome using the STAR aligner within Cell Ranger.

### Bioinformatics Analysis

After the Cell Ranger pipeline, all subsequent bioinformatics was performed by the authors. Cell Ranger outputs for each sample were initially read into R (v4.4.1) as a Single Cell Experiment using the ‘scran’ package (v1.32). Cells were filtered out if they met any of the following criteria: 1) <2000 or >40000 total reads; 2) <400 or >5500 expressed genes; 3) >15% mitochondrial reads as a percentage of total reads; or 4) >15% red blood cell reads (Hba-a1, Hba-a2, Hbb-bs, Hbb-bt, Hbb-bh1, Hbb-y, or Hba-x) as a percentage of total reads.

Log transformation and normalization were then performed using the logNormCounts() function in ‘scran’. The scran Single Cell Experiment objects were then converted to Seurat objects using the as.Seurat() function in the ‘Seurat’ package (v5.1.0). Both samples were then merged into a single Seurat object with the merge() function. The object was then split by genotype with the split() function, and transformed using the SCTransform() function. Dimensionality reduction was performed using the runPCA() function with default settings.

Sample integration was performed using the IntegrateLayers() function with the RPCAIntegration method. Nearest neighbors were calculated for the integrated data set using the FindNeighbors() function with 30 dimensions and a K parameter of 20. Clustering was then performed on the shared nearest neighbors (snn) using the FindClusters() function with a resolution of 0.2. Uniform manifold approximation and projections (UMAPs) were generated for the RPCA integrated data using the RunUMAP() function with 30 dimensions. Seurat object layers were then rejoined for further analysis using the JoinLayers() function.

Marker genes for each cell cluster were identified using the ‘scran’ findMarkers() function with binomial test type and only upregulated marker genes retrieved. The top 5 marker genes for each cluster were extracted and plotted using the ‘Seurat’ DotPlot() function. Cell clusters were then initially annotated from the top 20 marker genes in an unbiased method using the Annotation of Cell Types (ACT) online platform (http://xteam.xbio.top/ACT/)(PMID: 37924118), with Species = mouse, Tissue = Carotid artery segment. Unbiased results were then confirmed by literature search of marker genes and edited where appropriate.

The Seurat object was then subsetted for only EC using the subset() function, before repeating the ‘Seurat’ SCTransform, dimensionality reduction, integration, and clustering pipeline as above with the same settings. For clustering using the FindClusters() function, a resolution of 0.4 was used. The ‘scran’ findMarkers() function was then repeated as above to find marker genes for each EC population. The top 10 marker genes for each cluster were extracted and plotted using the ‘Seurat’ DotPlot() function. To generate a functional annotation for each cluster on a manageable subset of marker genes, pathway enrichment analysis was performed on the statistically significant EC cluster marker genes with a findMarkers() ‘Top’ rank < 201 with ‘clusterProfiler’ (v4.12.0) using the ‘org.Mm.eg.db’ (v3.19.1) and Gene Ontology (GO) biological process databases. GO terms with a padj < 0.05 following Benjamini-Hochberg multiple comparison adjustment were considered statistically significantly enriched and used to annotate cell clusters.

Differential gene expression (DGE) analysis was performed between EC-WT and EC-Cx43 KO groups separately for each identified EC cluster using the ‘Seurat’ FindMarkers() function with default settings. Given a small number of genes were identified as significant between genotypes using a padj > 0.05, subsequent pathway enrichment analysis was performed separately on upregulated and downregulated DEGs with an unadjusted p-value < 0.01 with ‘clusterProfiler’ and a log2(FC) < 0 and the top 10 pathways were plotted. Additionally, for the identified migratory cluster, all significantly enriched GO biological process terms containing the phrases ‘migration’, ‘erk’, ‘mapk’, or ‘protein kinase b’ were separately extracted and plotted. For the identified proliferative cluster, the same method was performed for the phrase ‘Cell cycle’. Gene ratio, gene count, and padj for the extracted pathways were plotted in R for visualization. Relative transcript expression was plotted for selected genes between groups in each EC cluster using the ‘Seurat’ VlnPlot() function.

## Statistical Analysis

Statistical tests were performed using GraphPad Prism 10 Version 10.4.1. Outliers were identified using Grubbs outlier test (alpha=0.05). 1-way or 2-way multiple comparison ANOVA followed by Dunnett post-test was used for comparisons between >2 groups, and T-test was used for comparisons of 2 treatment groups. A minimum of N=3 was used for all statistical analyses, n values for each experiment are stated in the figure legends. P values are shown for all statistical analyses, a P value of 0.05 is significant. Error bars represent mean ± SEM.

## Results

### Vascular Injury Upregulates Endothelial Cx43 Gap Junctions

Cx43 expression has been documented in large artery endothelial cells (ECs) experiencing heightened levels of shear-induced stress but was not previously thought to be present outside regions of vascular branching (27). To test the effect of injury-induced stress on EC Cx43 expression, we analyzed publicly available bulk RNA-seq data published by McDonald et al. (GEO: GSE115618) (20). In their study, endothelial injury was induced by applying a vascular clamp to a surgically exposed mouse aorta for 1 minute. After releasing the clamp, the mice recovered for either 3 or 48 hr. An adjacent area of the aorta served as a no-injury control. RNA lysis buffer was flushed through the desired portion of the vessel to collect the endothelium. The previous study demonstrated that connexins known to be expressed in large artery endothelium, GJA4 (Cx37) and GJA5 (Cx40), had high transcript levels in the aorta and that Cx37 expression was transiently upregulated at 3 hr post-injury which returned to uninjured levels by 48 hr. No change in in Cx40 expression was observed. We performed a retrospective analysis of the data for Cx43 expression. Our data analysis revealed Cx43 to be at undetectable levels in control tissues, but was significantly upregulated at both 3 hr and 48 hr timepoints (**Fig. 1A**), coinciding with reported timepoints of EC proliferation and migration responses (20).

**Figure 1:**
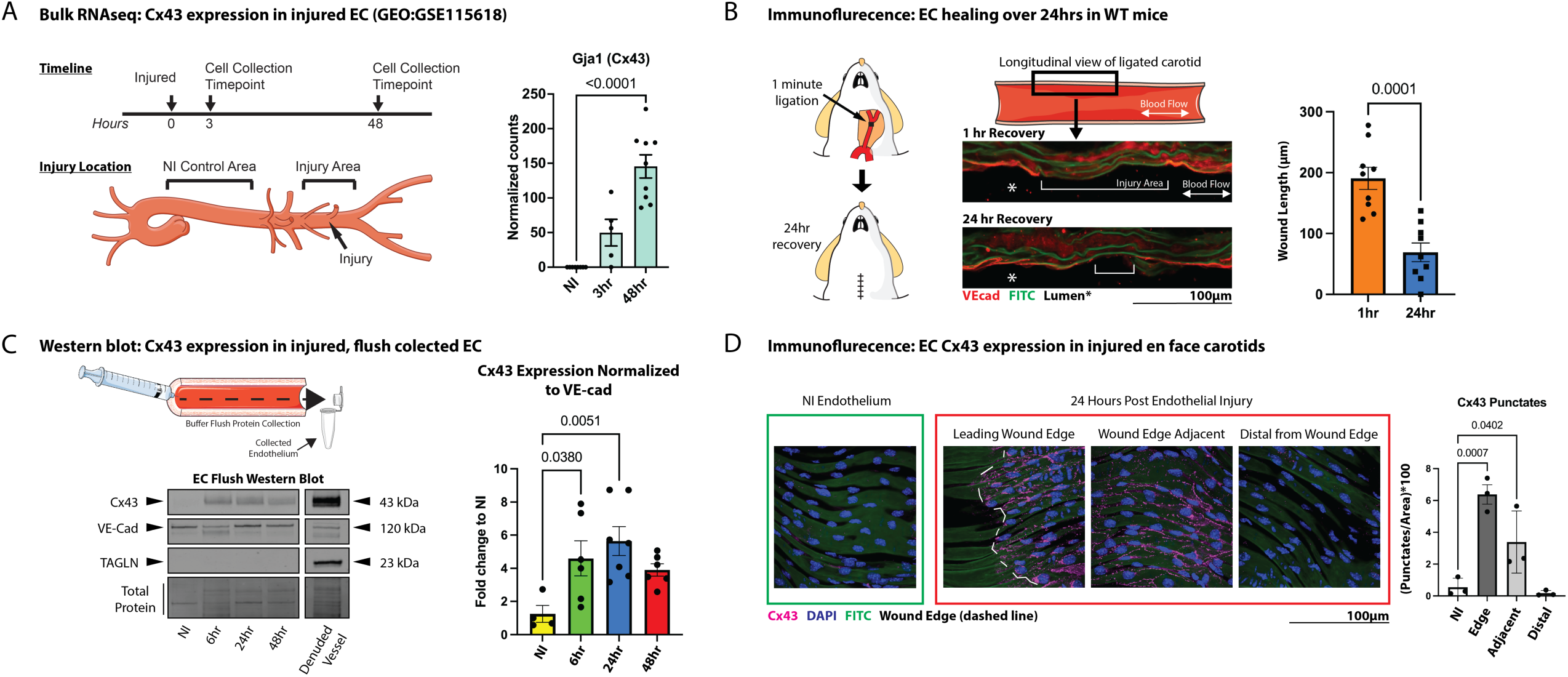
Endothelial injury upregulates Cx43 expression. Data adapted from McDonald et al. (GEO: GSE115618). Graphs show bulk RNA sequencing data from flush collected EC after one minute of vascular clamp injury followed by 3 or 48 hour recovery. Schematic shows injury and collection timepoints as well as location of injured EC and EC used for no injury controls (**A**). Schematic shows location of carotid artery ligation EC injury model. IF images show longitudinally sectioned, injured carotid arteries at 1 and 24hr post-injury timepoints (n=9) (**B**). Western blot of NI and injured flushed carotid artery EC allowed to recover for 6, 24, and 48hr. EC denuded vessels included as control (n=4-7) (**C**). IF detection of Cx43 expression in en face NI and EC injured carotid artery EC after 24hr recovery (**D**).

To test the role of Cx43 in EC repair *in vivo*, we developed a model of EC injury using the mouse carotid artery (42). Carotid ligation was chosen over aortic clamping due to ease of access and reduced surgical invasiveness. We found that vessel clamps used in aortic surgeries did not induce a clear section of injured EC as previously demonstrated in aortas (data not shown). One-minute carotid ligations resulted in a 190.6 ± 18.3 μm area of EC denudation after 1 hr of recovery, which was reduced to 69 ± 15.4μm after 24 hr of recovery. Vessel injury was visualized by immunofluorescence using the EC-specific marker VE-cad in longitudinally sectioned and en face carotids (**Fig. 1B, Supplemental Fig. 1 and 2**). To assess Cx43 expression at the protein level, we flushed protein lysis buffer through carotids that had undergone no injury (NI) or injury followed by 6, 24, or 48 hr of recovery. Western blotting demonstrated Cx43 expression along with VE-cad to confirm the presence of ECs and a lack of transgelin (TAGLN) contamination by smooth muscle cells. Flushed, EC-denuded vessels served as a TAGLN-positive control. Our data highlight a lack of detectable EC Cx43 in NI vessels followed by significant increases in EC Cx43 expression as early as 6 hr after injury, with expression peaking at 24 hr, just prior to wound closure (**Fig. 1C, Supplemental Fig. 3**.

To capture spatial localization of EC Cx43 expression after injury, we performed immunofluorescent staining on en-face carotids 24 hr post-injury. We found significant increases in punctate Cx43 expression at the leading edge of the EC wound and in cells adjacent (up to 500 µm) from the wound edge (**Fig. 1D, Supplemental Fig. 4**). In contrast, almost no Cx43 expression was detected in the uninjured endothelium or in ECs at sites distal (approximately >500 µm) from the wound.

### Cx43 Knockout Delays Endothelial Wound Closure

To assess the role of injury-induced EC Cx43 upregulation in wound healing, we bred mice expressing Cre recombinase specifically in EC (CDH5^(PAC)CreERT2^) mice with Cx43^fl/fl^ mice to generate tamoxifen-inducible EC-Cx43 KO mice and used Cre-negative Cx43fl/fl (EC-WT) littermate controls (**Fig. 2A**). Genotyping confirmed the EC-Cx43 KO genotype with both EC-Cx43 KO and EC-WT control mice containing Cx43 flox sites (**Fig. 2B and Supplemental Fig. 5**). Primers were designed to detect the deletion of Cx43 based on previous publication (76). Bands for Cx43 KO were detected in EC-Cx43 KO but not EC-WT aortic endothelium from tamoxifen-exposed mice (**Fig. 2B and Supplemental Fig. 5**). Immunofluorescent staining of Cx43 in injured carotid arteries from tamoxifen-exposed EC-Cx43 KO mice and EC-WT controls revealed loss of Cx43 expression only in the endothelium of EC-Cx43 KO mice (**Fig. 2C and Supplemental Fig.6**). One-minute carotid ligation surgeries in mice followed by 24 hr recovery was performed in EC-Cx43 KO and EC-WT mice. Data demonstrate significant delays in the rate of EC healing in EC-Cx43 KO (wound area 159.4 ± 20.2 μm) mice compared to EC-WT controls (wound area 83.8 ± 11.8 μm) (**Fig. 2D and Supplemental Fig. 7**).

**Figure 2:**
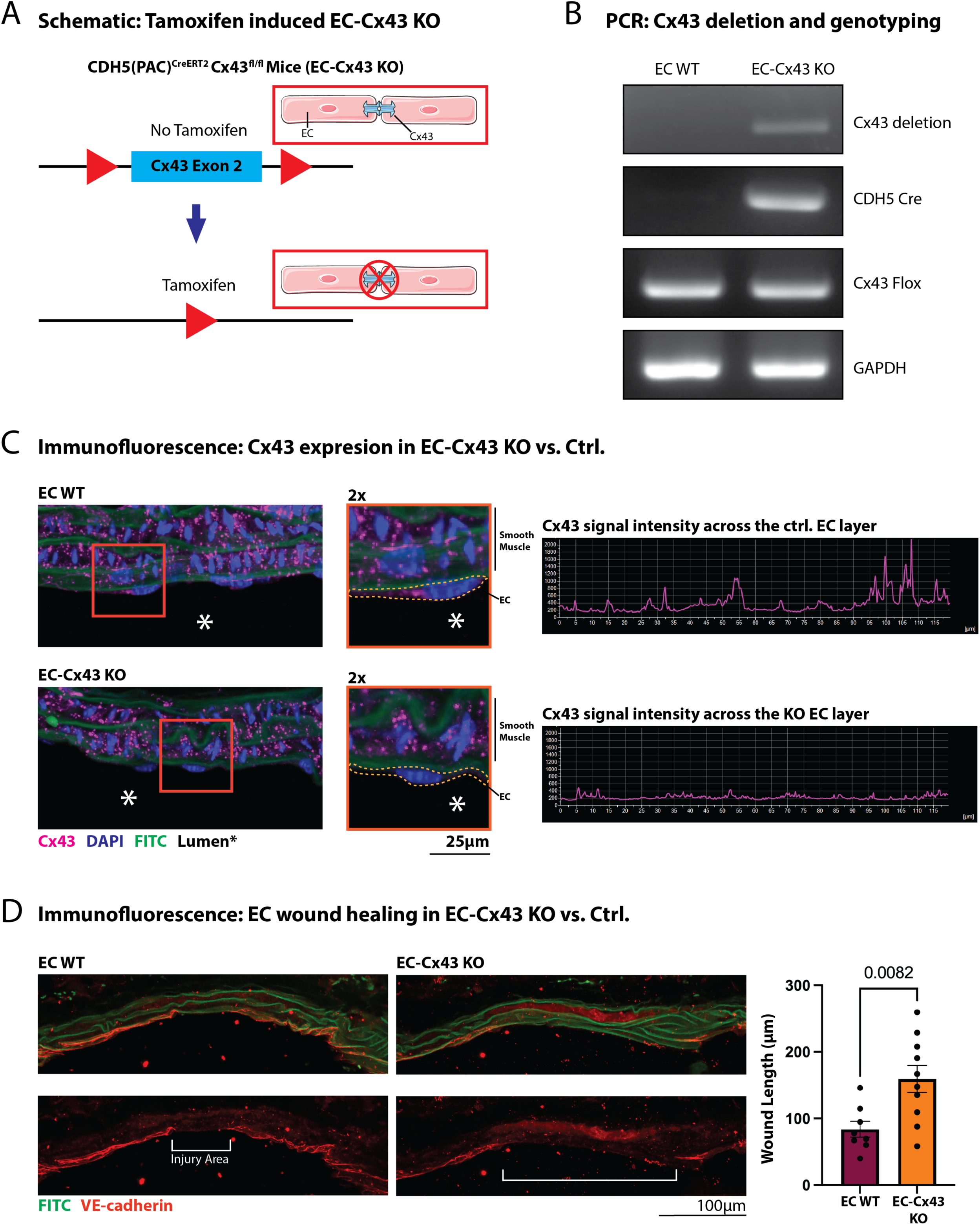
EC healing delayed in EC Cx43 KO mice. Schematic of tamoxifen inducible EC -Cx43 KO mice (CDH5(PAC)^CreERT2^, Cx43^fl/fl^) (**A**). PCR gel demonstrating expression of tamoxifen induced Cx43 deletion in Cx43^fl/fl^, Cre+ and Cre-control mouse EC (n=6) (**B**). IF detection of EC Cx43 in longitudinally sectioned carotids 24hr after injury in EC-Cx43 KO (Cre+) vs. EC WT (Cre-) control mice (n=3) (**C**). IF detection of EC (VE-cad) wound length 24hr after EC injury in EC-Cx43 KO vs control mice (n=7-8) (**D**).

### Single-Cell Sequencing Reveals Distinct Cell Populations in Whole EC-Cx43 EC KO vs. WT Injured Carotids

Cx43 expression after vascular injury is localized to EC that are in close proximity to the wound which are actively involved in wound response and regeneration. As such, bulk RNA sequencing approaches would provide only limited information on changes in specific populations of EC. Therefore, to assess the transcriptional response of Cx43-KO in regenerating EC, single-cell RNA sequencing was performed commercially by Novogene on EC-enriched carotid single-cell suspensions from injured EC-WT and EC-Cx43 KO at 18 hr post-injury (**Fig. 3A**). An 18 hr timepoint was selected to capture both the proliferative and migratory aspect of EC regeneration based on previously published healing timelines in large mouse arteries (20). Following sequencing, the EC-WT sample contained 7298 cells and the CDH5pos group contained 8315 cells. An average read depth of ∼65,000 raw reads per cell was achieved, which equated to a median of ∼4500 unique molecular identifier (UMI) counts that mapped to a median of ∼2000 genes per cell. After excluding poor quality cells and potential doublets, we analyzed 10,829 cells across 2 samples (**Supplemental Fig. 8**).

**Figure 3:**
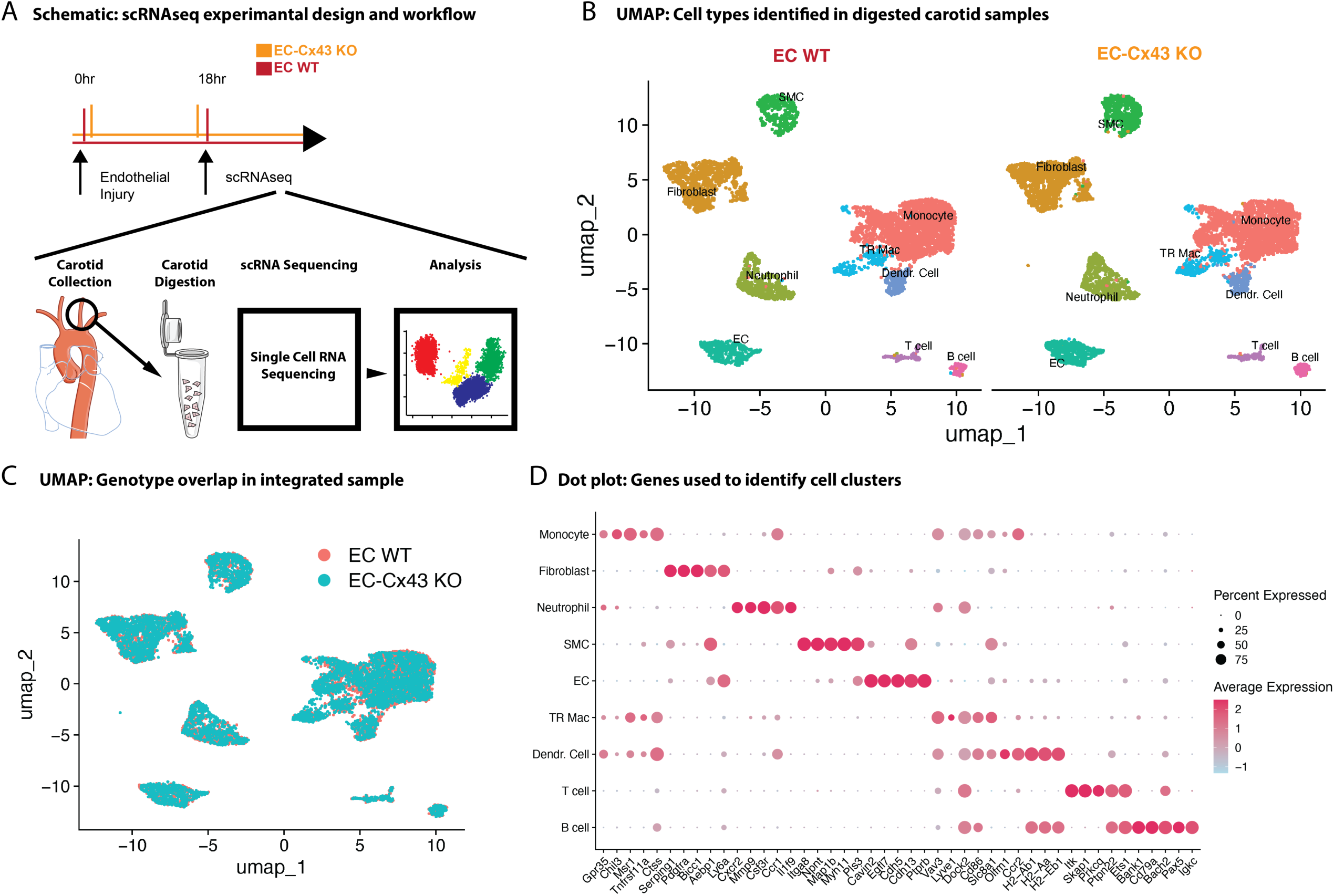
Endothelial populations present in injured carotid scRNAseq. Schematic of 18 hr post-injury carotid collection timepoint and enzymatic digestion for Novogene scRNAseq analysis in EC-WT control vs EC-Cx43 KO mice (**A**). Unilateral Manifold Approximation and Projection (UMAP) plot showing identified cell populations. Endothelial cell (EC), monocyte, tissue resident macrophage (TR Mac), dendritic cell (Dendr. Cell), neutrophil, T cell, B cell, fibroblast, and smooth muscle cell (SMC) populations were found (**B**). UMAP showing spatial overlap of cell populations between EC-WT control vs EC-Cx43 KO mice samples (**C**). Dot-plot of the top 5 ranked marker genes specific to each cell population (**D**).

UMAP dimensionality reduction and clustering revealed clearly distinguished populations representing all expected cell types from vascular tissue, including EC, vascular smooth muscle, fibroblasts, and immune cells (**Fig. 3B,C and Supplemental Fig. 8**). Following integration of the two samples, no changes in overall cell populations were identified in EC-Cx43 KO compared to control. A distinct EC population was identified with the top 5 detected marker genes of Cavin2, Egfl7, Cdh5, Cdh13, and Ptprb (**Fig. 3D**).

### EC-Cx43 KO Inhibits Endothelial Migration and Proliferation After Injury

As our initial data suggest that distinct subpopulations of EC in close proximity to the wound express Cx43, we extracted the subset of EC from the total scRNAseq dataset for further analysis and performed separate dimensionality reduction (**Figure 4A,B**). Our data contained 794 ECs which clustered into 4 subpopulations (**Figure 4B**). Pathway enrichment analysis using the Gene Ontology (GO) biological process database on identified marker genes from each EC subpopulation revealed that cluster 1 had significant enrichment of migratory pathways, cluster 2 had significant enrichment of cell cycle pathways and was the only cluster with strong expression of the cell cycle gene Ccnd1, and cluster 3 had significant enrichment of pathways involved in immune cell interactions. Due to a lack of enrichment of distinctive pathways that would allow classification, and as the largest cluster, cluster 0 was designated as the quiescent EC population that would likely represent EC distal from the wound area (**Fig. 4C and Supplemental Fig. 9**). There was no difference in any subpopulation size as a percentage of total cells between genotypes (**Fig. 4B**).

**Figure 4:**
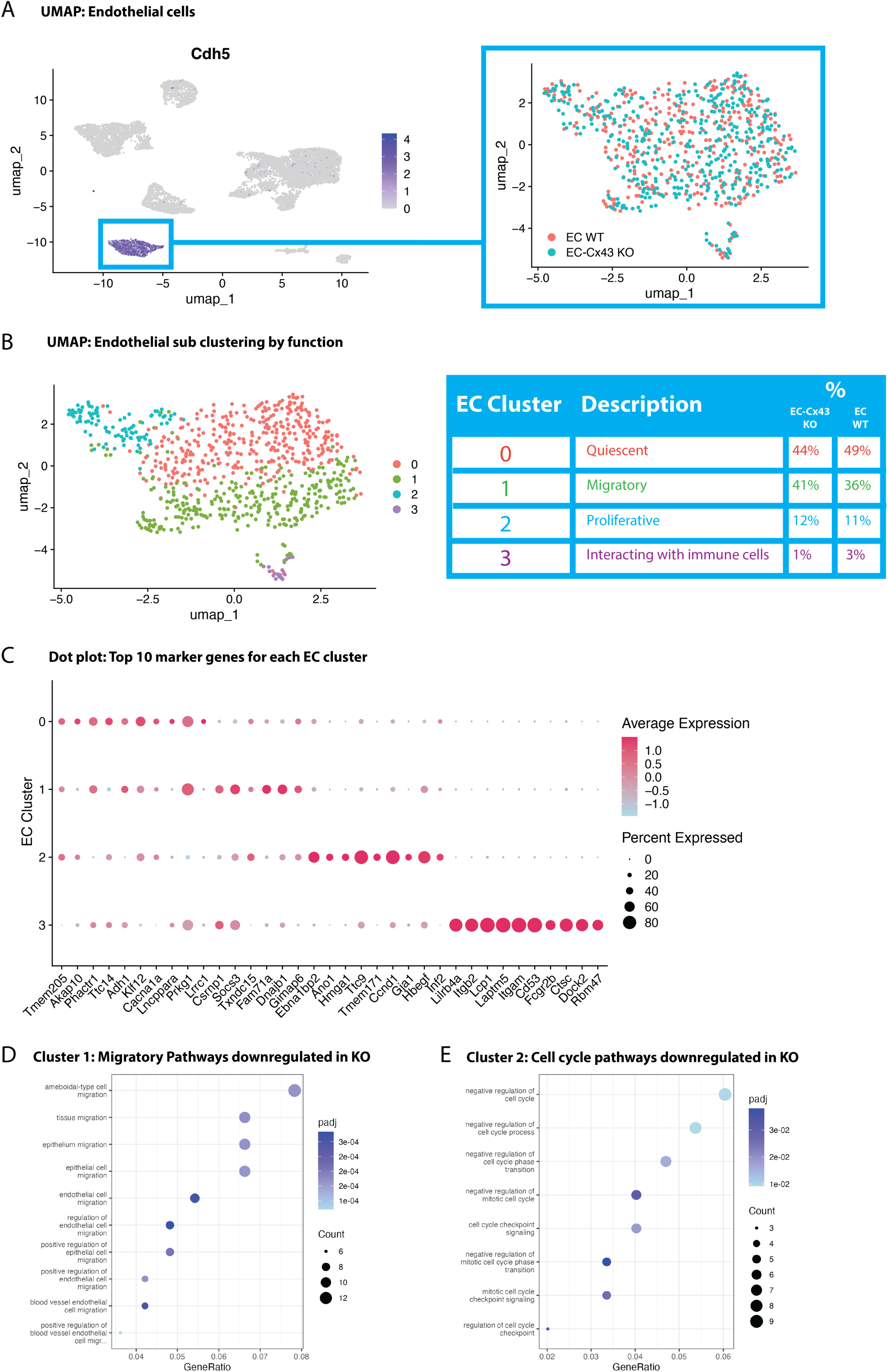
Cx43 KO suppresses markers of EC proliferation/migration. Feature plot of integrated cells colored by the endothelial cell (EC) marker CDH5 (left panel); and Unilateral Manifold Approximation and Projection (UMAP) plot of the integrated CDH5^+^ EC subset colored by genotype (right panel; **A**). Clustered UMAP of integrated EC with table indicating annotation and total cells in each cluster as a percentage of all EC per genotype (**B**). Dot-plot of the top 5 ranked marker genes specific to each EC cluster (**C**). Migratory pathways from cluster 1 (**D**) and cell cycle pathways from cluster 2 (**E**) extracted from Gene Ontology (GO) biological process enrichment analysis of downregulated genes in EC-Cx43 KO cells compared to EC-WT control endothelial cells.

As proliferative and migratory EC subpopulations are likely actively involved in EC wound healing, we assessed the role of Cx43 knockout on the transcriptional response to injury in these groups. Given the low number of cells, the cluster designated as immune interacting was not analyzed further. As expected in a knockout model, few meaningfully enriched GO pathways were observed in the list of significantly upregulated genes (**Supplemental Fig. 10**). GO biological process pathway enrichment analysis of genes significantly downregulated in EC-Cx43 KO compared to control cells at the unadjusted p<0.01 level revealed significant downregulation of migratory pathways, such as endothelial and epithelial cell migration, in the migratory EC cluster (**Fig. 4D and Supplemental Fig. 11**), and proliferative pathways, related to cell cycle regulation and transition, in the proliferative EC cluster (**Fig. 4E**) in response to EC-Cx43 KO.

### Phosphorylation Regulated Gap Junction Signaling Influences EC Wound Healing

Cx43 is highly regulated by phosphorylation resulting in altered gap junction functionality and cell signaling (81). We used our scRNAseq data to identify Cx43-dependent changes in pathways related to cell growth and division. Pathway enrichment analysis from EC-Cx43 KO downregulated DEGs identified downregulated protein kinase B (AKT) and ERK (MAPK) signaling in migratory cluster 1 (**Fig. 5A**). Both AKT and MAPK can phosphorylate the Cx43 c-terminus which can alter gap junction functions (82, 83). To investigate whether phosphorylation of the Cx43 c-terminus contributes to Cx43 control of EC wound healing functions, we performed endothelial injury surgeries in the carotids of genetically modified mice with knock-in phospho-null alanine mutations at Cx43-PKA/ AKT, Cx43^S255/262/279/282A^, and Cx43^S368A^ sites respectively (**Fig. 5B**). Measurements of EC injury wound areas at 24 hr post injury show a significant delay in wound closure in in Cx43^S255/262/279/282A^ mice(154.7 ± 22.2 µm) mice compared to C57BL6-J mice (69 ± 15.4 µm) (**Fig. 5C and Supplemental Fig. 12**). No significant changes in EC healing rate were identified in Cx43^S365/373A^ and Cx43^S368A^ mice.

**Figure 5:**
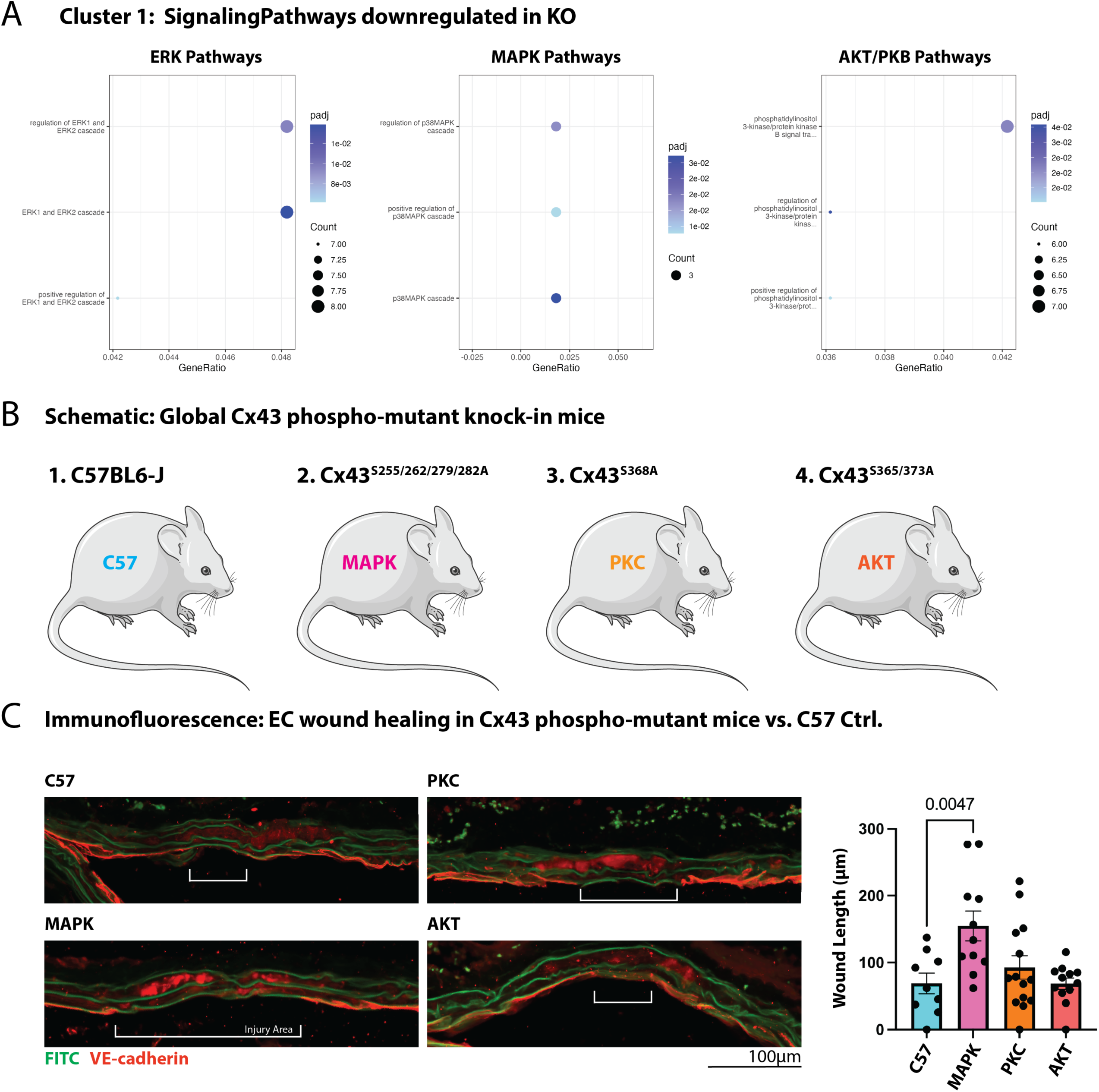
Cx43 regulation of EC healing dependent on Cx43-MAPK phosphorylation. ERK (left panel), MAPK (middle panel), and AKT/PKD (right panel) pathways extracted from Gene Ontology (GO) biological process enrichment analysis of downregulated genes in EC-Cx43 KO cells compared to EC-WT control cluster 1 (migratory) endothelial cells **(A)**. Schematics showing mouse phospho-null genotypes used for EC injury experiments **(B)**. IF images show longitudinally sectioned, injured carotid arteries at 24hr post-injury in C57, MAPK, PKC, and AKT phospho-null mice. To avoid unnecessary duplication in mice, C57 24 hr EC injury data is from Fig. 1B. White brackets indicate the length of EC injury (n=9-13) **(C)**.

## Discussion

Endothelial cell (EC) regeneration is crucial for vascular healing and long-term patency of vessels following surgical intervention. Delayed EC healing significantly contributes to vascular vulnerability to thrombosis immediately after surgery. Furthermore, prolonged disruptions to the endothelial layer can result in inflammation, the reemergence of atherosclerotic disease, or the pathological phenotypic switching of vascular smooth muscle cells, ultimately leading to neointima formation. Understanding the factors that regulate these healing processes is essential for the development of future vascular therapeutics. In this study, we identify the gap junction protein Cx43 as a key regulator of EC regeneration and vascular wound closure. We demonstrate that Cx43 expression is specifically upregulated in EC responsible for healing following injury, and that preventing Cx43 upregulation via genetic knockout delays the healing process. We provide data that highlights pathways involved in Cx43 control over EC proliferation and migration, and identify that phosphorylation of Cx43 at known phosphorylation sites targeted by mitogen-activated protein kinases (MAPK) is a key mechanism for this control. These results suggest that injury induced Cx43 expression and gap junction functionality are important for normal wound healing in the vasculature.

In 2020, the ISCHEMIA trial revealed that patients who undergo invasive procedures to correct coronary artery disease do not have superior outcomes in terms of cardiovascular events like heart failure, cardiac arrest, or even death from cardiovascular causes compared to patients who undergo medical therapy alone (4). These findings were driven by adverse events occurring at early time points after invasive procedures were performed (4). Patients undergoing invasive procedures who made it past these early initial time points experienced improved quality of life measures and an improved likelihood of survival (4). As such, it is essential to develop safer surgical practices to ensure patients benefit from surgery. Endothelial damage is intrinsically linked to early failure of these surgical interventions, and endothelial healing after surgery should be a primary concern.

Our previous investigation revealed extensive endothelial damage in saphenous veins prepared for coronary artery bypass grafting (9). This finding has been demonstrated in multiple other studies suggesting that there is an unavoidable loss of EC coverage using current best practices in surgical preparations (15, 16). In PCI, stents also cause significant damage to the endothelium, and their physical barrier and anti-proliferative drug coatings impair the endothelial healing process. In rabbits, the placement of a bare metal stent resulted in delayed endothelial healing compared to balloon angioplasty, while the placement of a standard mTOR inhibiting drug-eluting stent delayed healing further and resulted in transcriptional changes indicative of activated, unhealthy EC (84). Current strategies for mitigating the effects of delayed EC healing include the use of dual antiplatelet therapy, sometimes for over a year after stent placement (85). This strategy itself can be dangerous, as it increases the risk of major bleeding (85). All these factors together reveal the necessity of better methods of coronary intervention that account for EC health and repair.

The timeline of EC healing and the relative contributions of proliferation and migration were previously demonstrated in mice by McDonald et al., who showed that migration occurs prior to the onset of proliferation, with proliferation peaking around the time of wound closure and continuing into later time points (20). Our reanalysis of their bulk RNA-seq data from injured, flushed mouse aorta revealed that Cx43 begins to be expressed at the RNA level in the arterial lumen adjacent cells. Early studies identified Cx43 expression specific to large artery EC exposed to turbulent flow, concluding that Cx43 expression in these cells is a specific response to mechanical stress (27). In our model of carotid artery EC wounding, Cx43 protein expression appears in response to mechanical injury, with expression peaking near the time of wound closure and gradually tapering off at later stages. Immunofluorescent staining revealed strong Cx43 punctate membrane localization at the wound leading edge EC and in adjacent EC, but not at distal regions. This suggests that newly expressed Cx43 forms functional gap junctions and facilitates intercellular communication that may coordinate migratory and proliferative responses from cell to cell. Our observations that Cx43 proteins are not expressed at distal sites to the wound edge, suggest Cx43 plays a specific function in cells associated during wound healing in vivo.

Connexins have long been associated with cellular wound healing functions. In skin, Cx43 undergoes dynamic changes in response to wounding. For example, in epidermal keratinocytes *in vivo,* Cx43 is downregulated in response to wounding and Cx43 knockdown via the application of topical antisense oligonucleotides improves wound healing and reduces scarring (86–88). However, upregulation of Cx43 in the dermis has also been observed in areas undergoing heightened levels of proliferation in response to wounding, calling the exact role of Cx43 in wound healing into question (89). The mechanisms underlying these changes are complex due to the diverse cell types present in the skin, including capillary ECs, fibroblasts, and immune cells that express Cx43 at the wound site. In the present study, we find that EC-Cx43 KO delays wound healing in vivo, demonstrating that Cx43 upregulation specifically occurs as a response to EC injury and promotes wound healing in this context.

A key factor in epithelial wound healing is epithelial to mesenchymal transition. Its endothelial counterpart, endothelial to mesenchymal transition (EndMT), is a process in which EC markers associated with endothelial identity, such as VE-cadherin, are downregulated, while mesenchymal markers, including α-SMA, are upregulated (90). After undergoing EndMT, EC disassociate from surrounding cells and begin to migrate, potentially serving as a protective mechanism to expedite wound closure (91). However, ECs that have undergone EndMT may lose essential functions, such as regulating platelet aggregation, preventing thrombosis, and maintaining vascular barrier integrity through tight and adherens junctions. Partial EndMT, characterized by the expression of CD45 along with other EndMT markers such as α-SMA, has been reported in response to vascular damage (92). Additionally, Cx43 has been implicated in promoting the progression of EndMT in cultured and corneal EC (93, 94). Given our observations that Cx43 is upregulated in large artery ECs after injury and that Cx43 knockout (KO) impairs wound healing, we used single-cell RNA sequencing (scRNA-seq) to investigate the influence of Cx43 on endothelial identity, proliferation, and migration. From our scRNA-seq dataset, we identified distinct subpopulations of ECs undergoing proliferation and migration in response to injury, which likely correspond to those EC that are closest to the site of injury. Knockout of EC-Cx43 reduced the enrichment of pathways involved in both proliferation and migration in their respective clusters. Further interrogation of DEGs revealed the downregulation of genes associated with mesenchymal transition in EC-Cx43 KO mice (Itga3, Plk3, Ilk) (95–98). Based on these findings it is possible that injured ECs rely on Cx43-dependent mesenchymal transition to facilitate efficient wound healing. Interestingly, we did not observe an increase in markers of advanced EndMT such as α-SMA and vimentin, in our scRNAseq EC data. This lack of increase may be attributed to the 18 hr time point being insufficient to allow full EndMT to occur. Although peak proliferation and migration are likely to occur before complete EC healing takes place around our 18hr timepoint, both proliferation, and migration are likely to continue even past complete wound closure (20). We hypothesize that later time points beyond 24 hr might reveal more pronounced changes in EC identity and function, potentially including complete or partial EndMT.

To uncover the mechanism of delayed EC healing after Cx43 KO, we assessed Cx43-dependent changes in wound healing-related signaling pathways using our scRNAseq gene expression data. We found enrichment of both MAPK and AKT (protein kinase B) signaling pathways following analysis of genes downregulated by EC-Cx43 KO in the migratory EC cluster. Cx43 functionality is highly dependent on phosphorylation by kinases including MAPK, AKT, and PKC (78, 81). As such, we used Cx43^S255/262/279/282A^, Cx43^S368A^, and Cx43^S365/373A^ mice with phospho-null serine to alanine mutations at their respective phosphorylation sites to assess Cx43 phosphorylation-dependent changes in EC healing. EC healing in the Cx43^S255/262/279/282A^ phosphorylation-deficient mice was delayed to a similar degree to that of EC-Cx43 KO mice. In contrast, healing occurred at a normal rate in phosphorylation-deficient mutants for both Cx43^S368A^ and Cx43^S365/373A^. MAPK phosphorylation of Cx43 is associated with the final stages of the gap junction lifecycle, particularly the removal of gap junction channels from the cell membrane, and a downregulation of channel function (45, 53, 56, 99). This data suggest that MAPK phosphorylation mediated gap junction turnover is uniquely required for normal EC wound healing functions of Cx43. This agrees with our previous publication, which demonstrated that Cx43^S255/262/279/282A^ phosphorylation regulates phenotypic switching and subsequent proliferation in vascular smooth muscle cells (45). Cx43-PKC phosphorylation is associated with a decreased probability of gap junction channel opening and gap junction disassembly (46, 56). As such, Cx43^S368A^ mutant mice likely retain high numbers of functional gap junctions, explaining the maintained healing ability(100). In contrast to Cx43-MAPK and Cx43-PKC phosphorylation, Cx43-AKT phosphorylation occurs early in the gap junction lifecycle and precedes aggregation of gap junctions into larger gap junction plaques (53). As we do not see healing delays in these mice, it is possible that preventing Cx43-AKT phosphorylation does not significantly impair gap junction channel function and may explain why no delays in EC healing in the Cx43^S365/373A^ mice were found.

Our study reveals that through its regulation of migration and proliferation, Cx43 gap junction channels play a central role in restoring endothelial integrity. However, prolonged disruptions to EC identity, such as those induced by EndMT, may compromise vascular function and contribute to disease progression. One limitation of this study is that our model is only designed to assess wound healing in healthy EC and that the EC injuries used are not expected to produce long-term disease. Our data shows that Cx43 expression reduces after EC wound healing is complete. It is possible that in a model of chronic EC damage and vascular disease, Cx43 expression would remain high, and EndMT may be more pronounced, further contributing to long-term vascular dysfunction. Future research should also focus on understanding Cx43 in human EC healing and under stenting/bypass conditions. Our scRNAseq analysis provides high cell-cell resolution and is the first to show clear transcriptomic delineation between quiescent and proliferative/migratory EC in injured mouse carotid arteries. Despite this clear strength of the current study, limitations to this analysis exist. The physical size of the mouse carotid artery necessitates the pooling of multiple mice into a single sample, preventing the ability to analyze multiple independent replicates to enhance downstream functional gene analysis. Additionally, the reduced read depth achieved with current single-cell technology may mean that important changes between genotypes in this study are missed. Further research is required to fully characterize the downstream signaling events following EC injury injury-induced Cx43 upregulation and MAPK phosphorylation, resulting in EC phenotypic transition.

In conclusion, this study is the first to show that Cx43 protein expression is upregulated in large artery EC, which do not normally express Cx43, *in vivo* in response to injury. We demonstrate that both EC-specific Cx43 deletion and global Cx43^S255/262/279/282A^ mice display reduced healing. This is important because modulating Cx43 offers a promising therapeutic avenue for enhancing vascular healing while minimizing pathological outcomes to improve long-term vascular surgery success rates.

## Acknowledgements

We gratefully acknowledge the Histology resources and services provided by Dr. Clare Dennison, PhD at the FBRI Histology Core Facility at VTC. Servier medical art, licensed under CC BY 4.0 https://creativecommons.org/licenses/by/4.0/, was used to create figure schematics. The authors acknowledge the assistance of ChatGPT 4 for grammatical edits to enhance manuscript clarity coding assistance in R. This tool was used in a manner that does not conflict with APS ethical policies and the authors take full responsibility for the content.

## Data Availability Statement

Source data for this study is available upon request.

## Author contributions

MS, MR, BI, and SRJ conceived and designed research; MS, MR, KR, XL, CD, AB, BI, and SRJ performed experiments; MS, MR, KR, BI, and SRJ analyzed data; MS, MR, BI, and SRJ interpreted results of experiments; MS, MR, KR, and SRJ prepared figures; MS, MR, and SRJ drafted the manuscript; MS, MR, BI, and SRJ edited and revised manuscript; All authors approved final version of manuscript.

---

**Supplemental Fig. 1:**
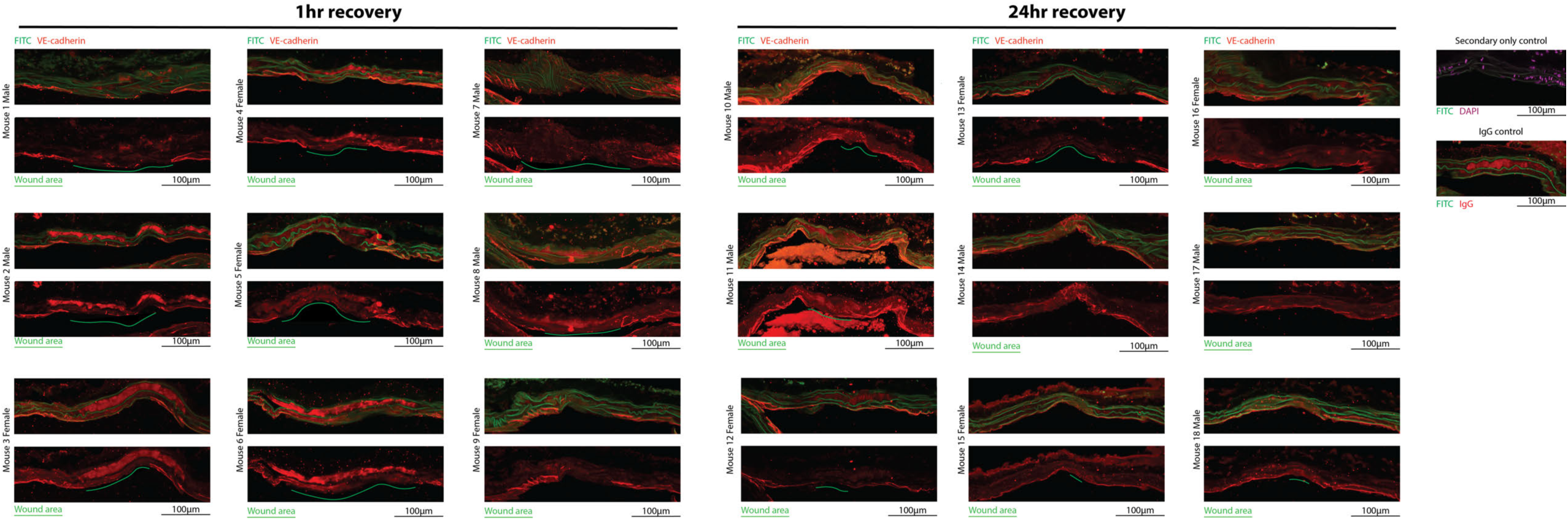
Immunofluorescent longitudinal view of C57 mouse carotids that underwent endothelial injury followed by 1 hour or 24 hour recovery. For each carotid, tissue autofluorescence and EC marker VE-cadherin are overlayed in the top image with VE-cadherin only in the bottom image. Green lines indicate the EC denuded area (n=9).

**Supplemental Fig. 2:**
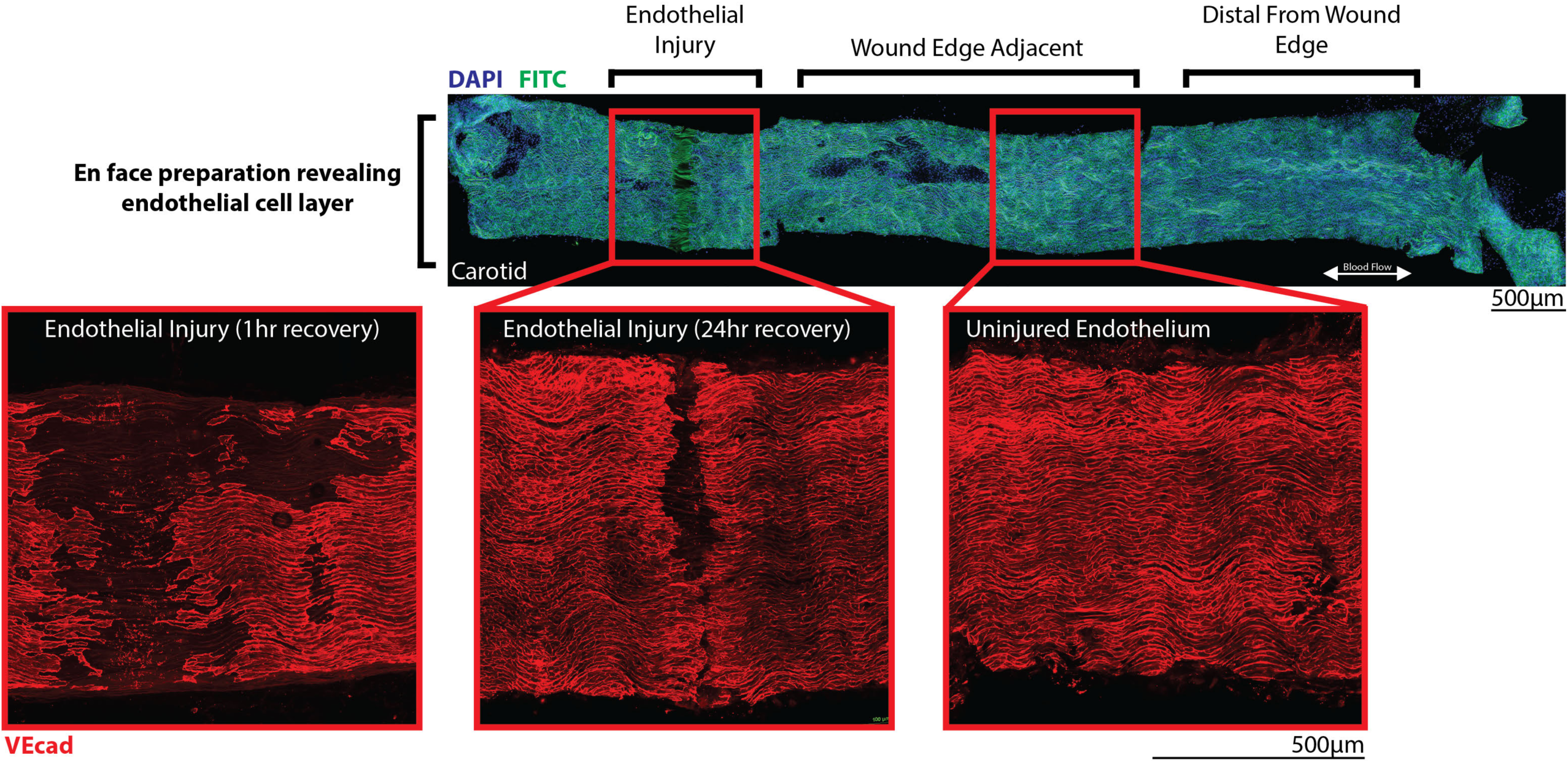
IF detection of VE-cadherin (EC) expression in en face EC injured carotid artery after 1 hour and 24hr recovery along with uninjured endothelium to show wound injury area.

**Supplemental Fig. 3:**
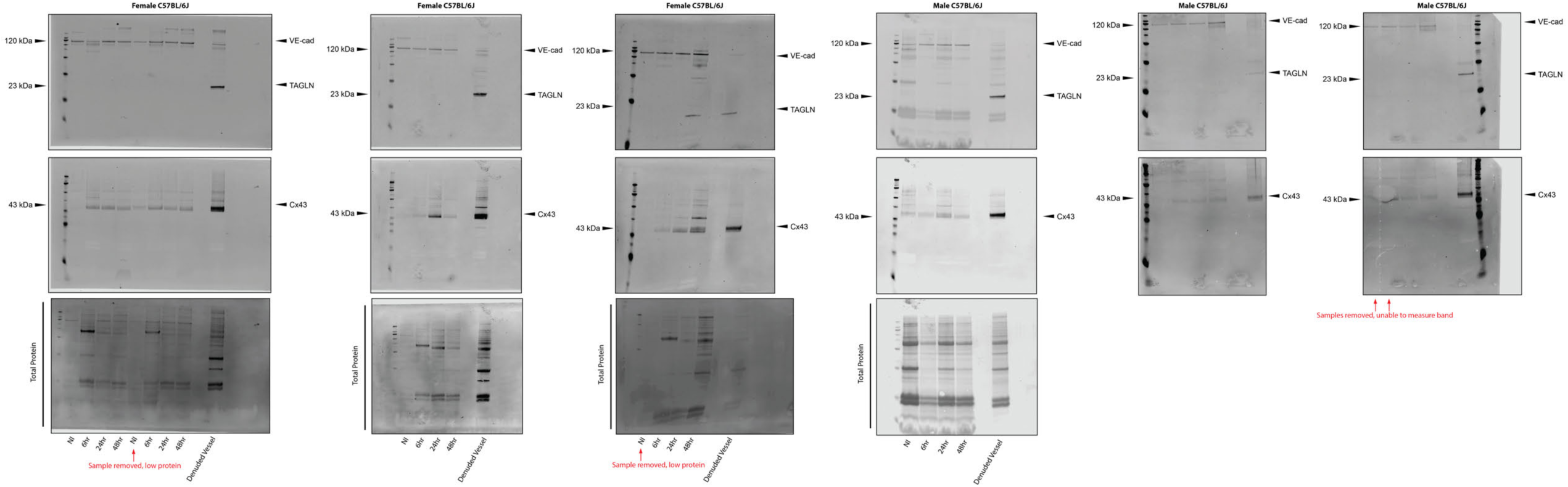
All EC flush western blots for no injury (NI), 6, 24, and 48 hour injury carotids with denuded vessel control (n=4-7). Antibodies are VE-cadherin (EC), TAGLN (smooth muscle), and Cx43.

**Supplemental Fig. 4:**
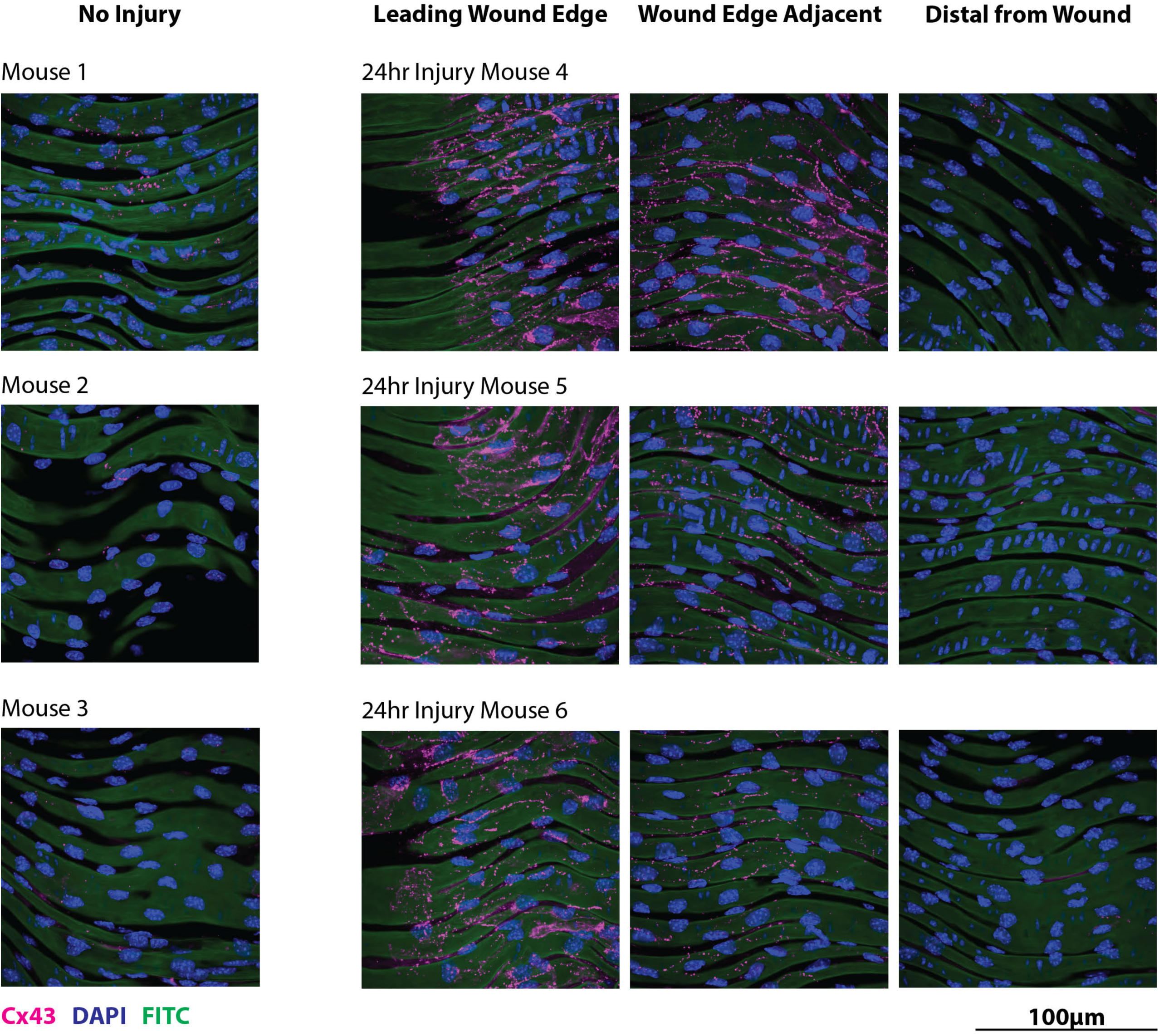
IF detection of Cx43 expression in en face NI and EC injured carotid artery EC after 24hr recovery. FITC shows tissue autofluorescence indicating the internal elastic lamina and DAPI staining identifies cell nuclei (n=3).

**Supplemental Fig. 5:**
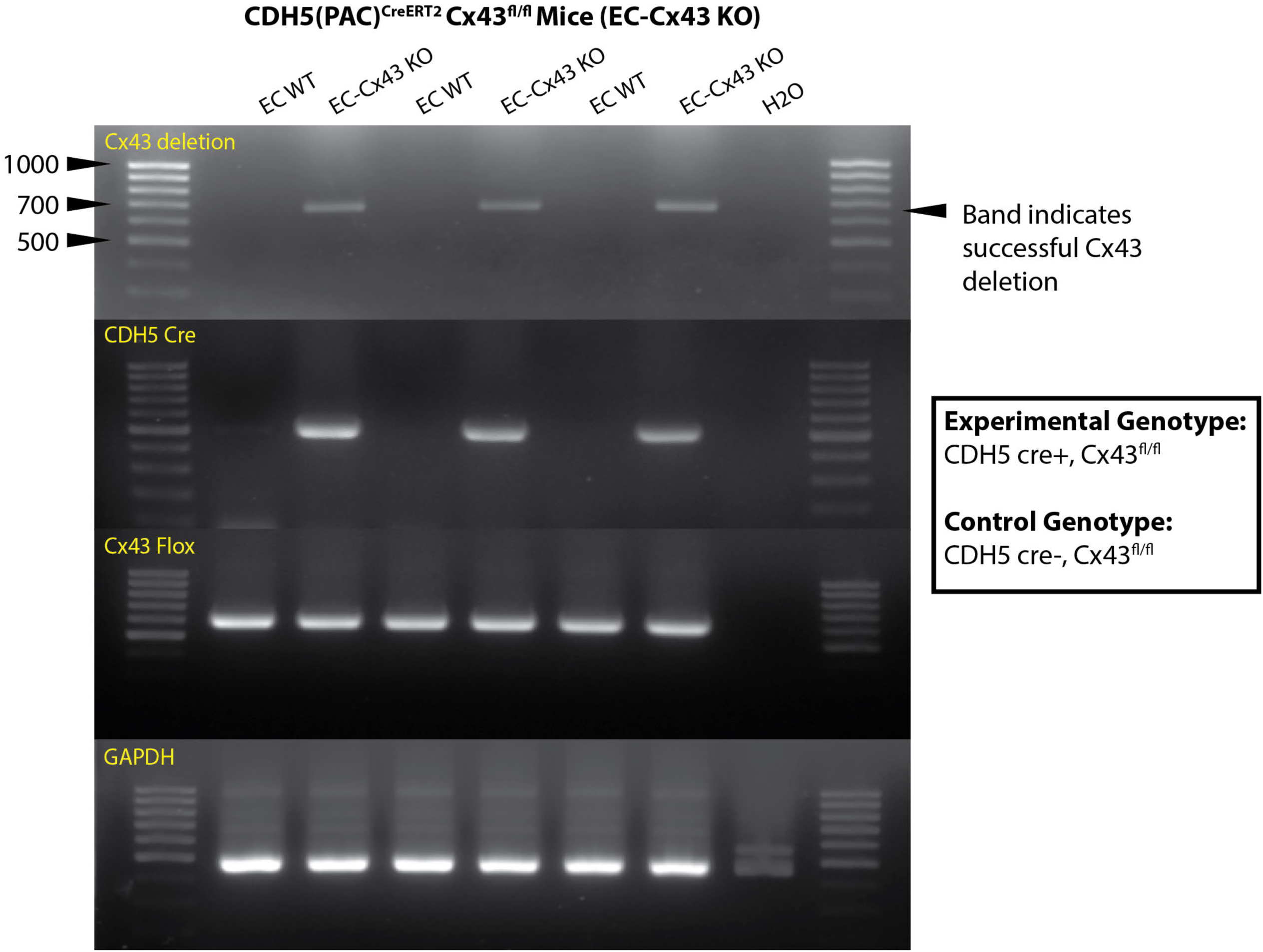
PCR gel demonstrating expression of tamoxifen induced Cx43 deletion in Cx43^fl/fl^, Cre+ and Cre-control mouse EC. Mouse genotypes confirmed using CDH5 Cre and Cx43 flox primers with GAPDH control (n=3).

**Supplemental Fig. 6:**
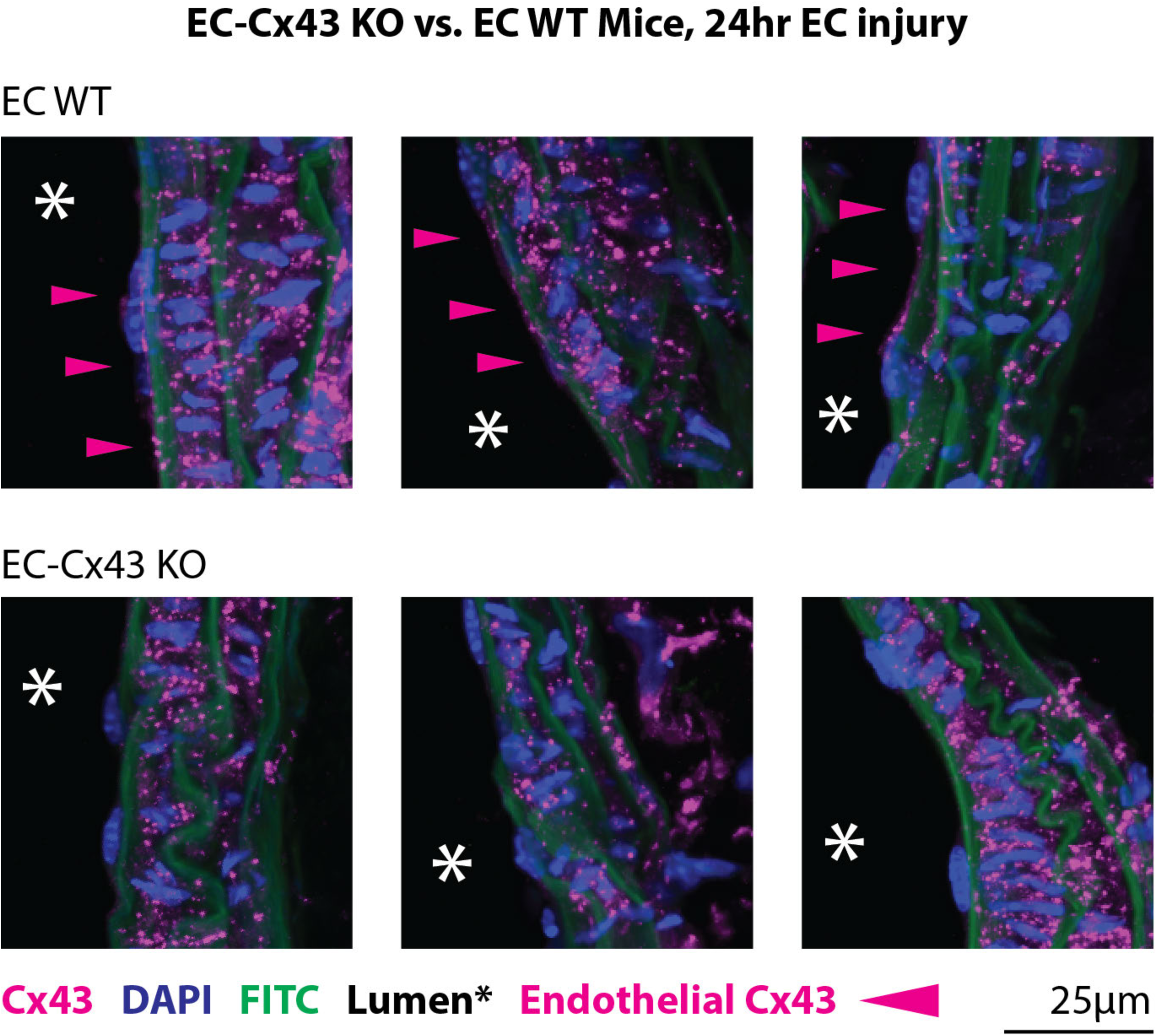
IF detection of EC Cx43 in longitudinally sectioned carotids 24hr after injury in EC-Cx43 KO vs. EC WT control mice. Pink arrows indicate endothelial Cx43 expression (n=3).

**Supplemental Fig. 7:**
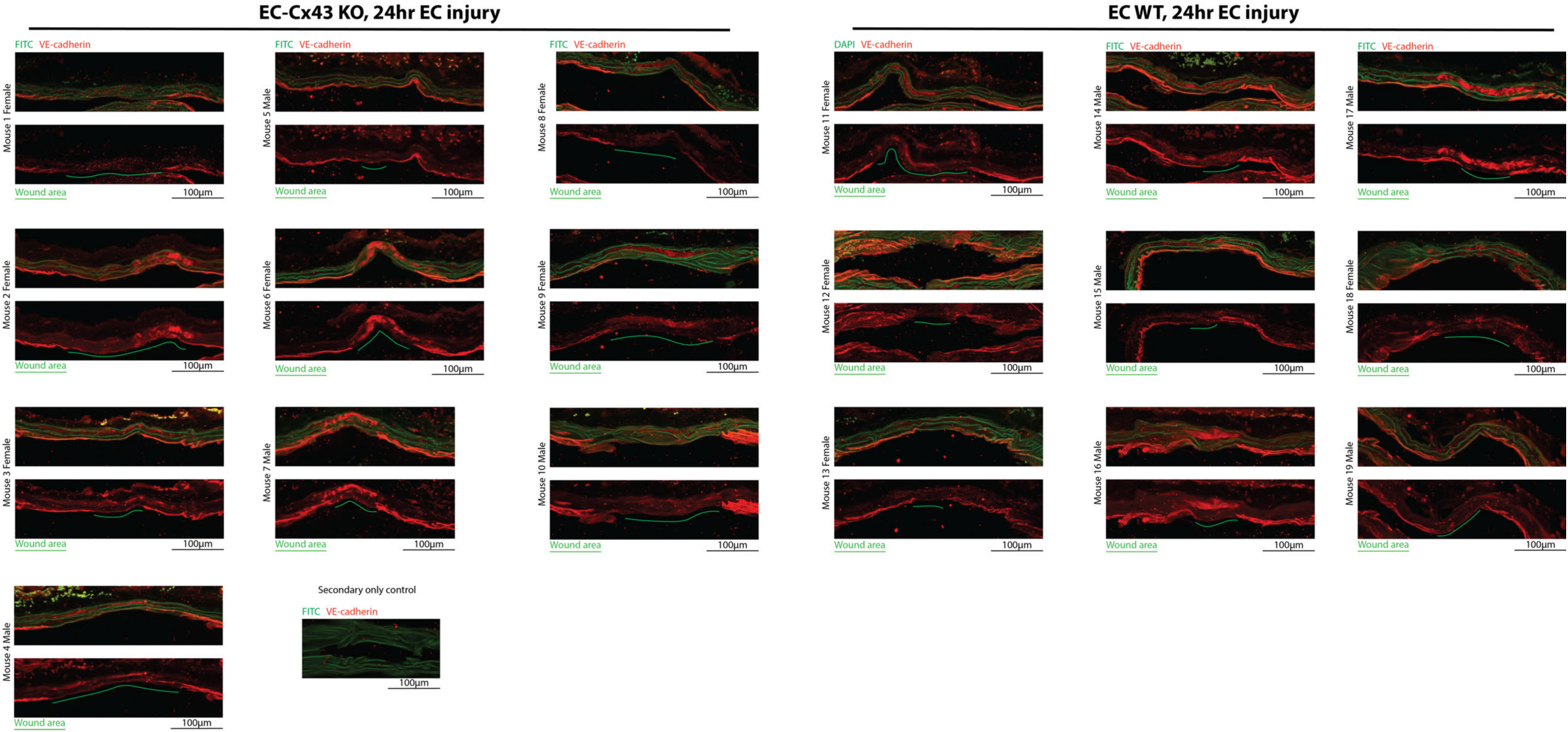
Immunofluorescent longitudinal view of EC-Cx43 KO and EC WT control mouse carotids that underwent endothelial injury followed by 24 hour recovery. For each carotid, tissue autofluorescence and EC marker VE-cadherin are overlayed in the top image with VE-cadherin only in the bottom image. Green lines indicate the EC denuded area (n=7-8).

**Supplemental Fig. 8:**
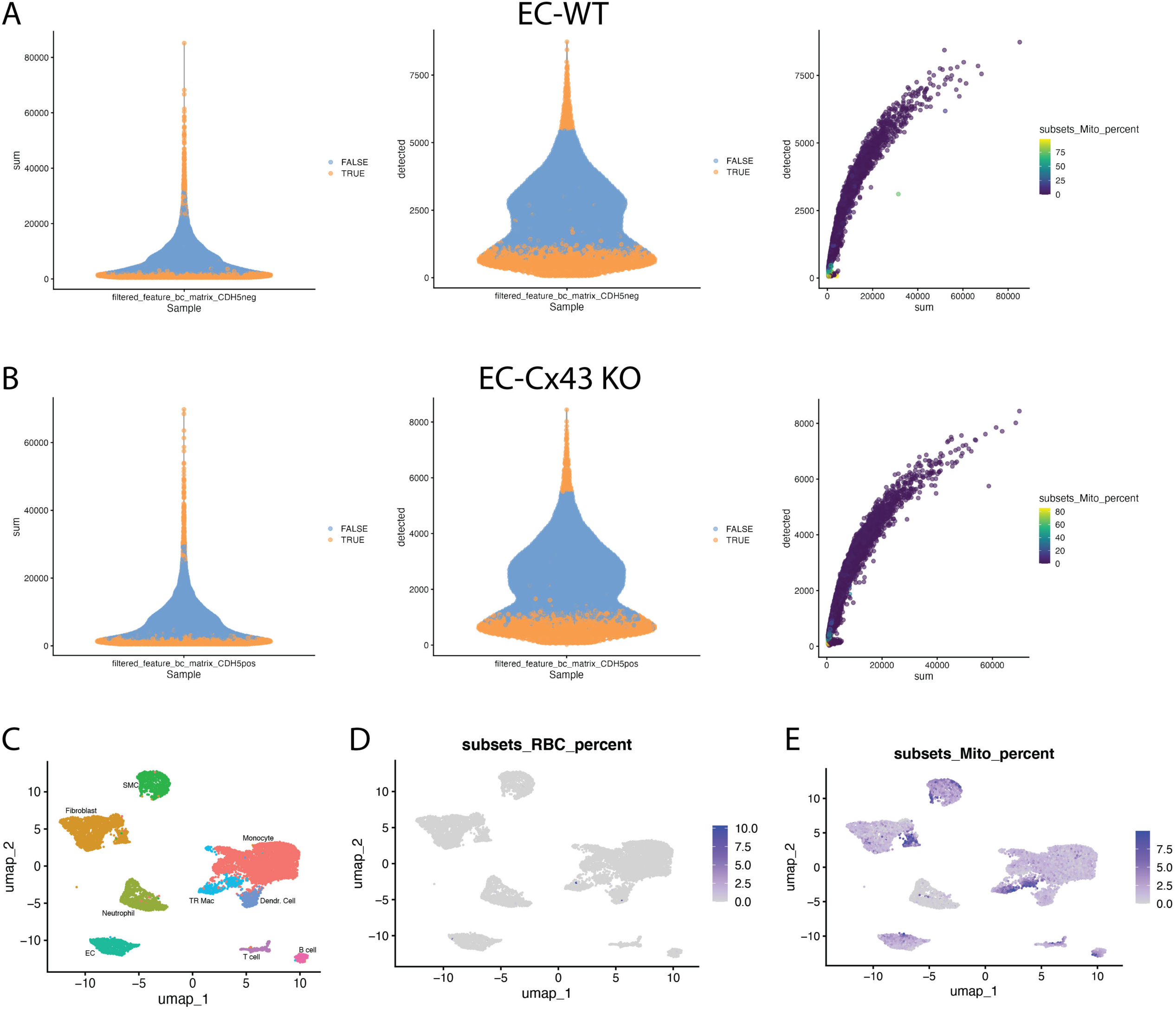
Mouse Carotid scRNAseq quality control analysis. Violin plots showing distribution of total reads (sum; left panel) and total genes detected (detected; middle panel) per cell with excluded low-quality cells shaded orange and cells included in the analysis shaded in blue, and scatter plots comparing total reads (sum) and total genes detected (detected) colored by total mitochondrial genes as a percentage of total reads (right panel) for EC-WT (**A**) and EC-Cx43 KO (**B**) mice. Unilateral Manifold Approximation and Projection (UMAP) of integrated cells from both EC WT and EC-Cx43 KO mice (**C**) Endothelial cell (EC), monocyte, tissue resident macrophage (TR Mac), dendritic cell (Dendr. Cell), neutrophil, T cell, B cell, fibroblast, and smooth muscle cell (SMC) populations were found. Feature plots colored by total red blood cell (RBC)-related genes as a percentage of total reads (**D**) and total mitochondrial genes as a percentage of total reads (**E**) in integrated cells.

**Supplemental Fig. 9:**
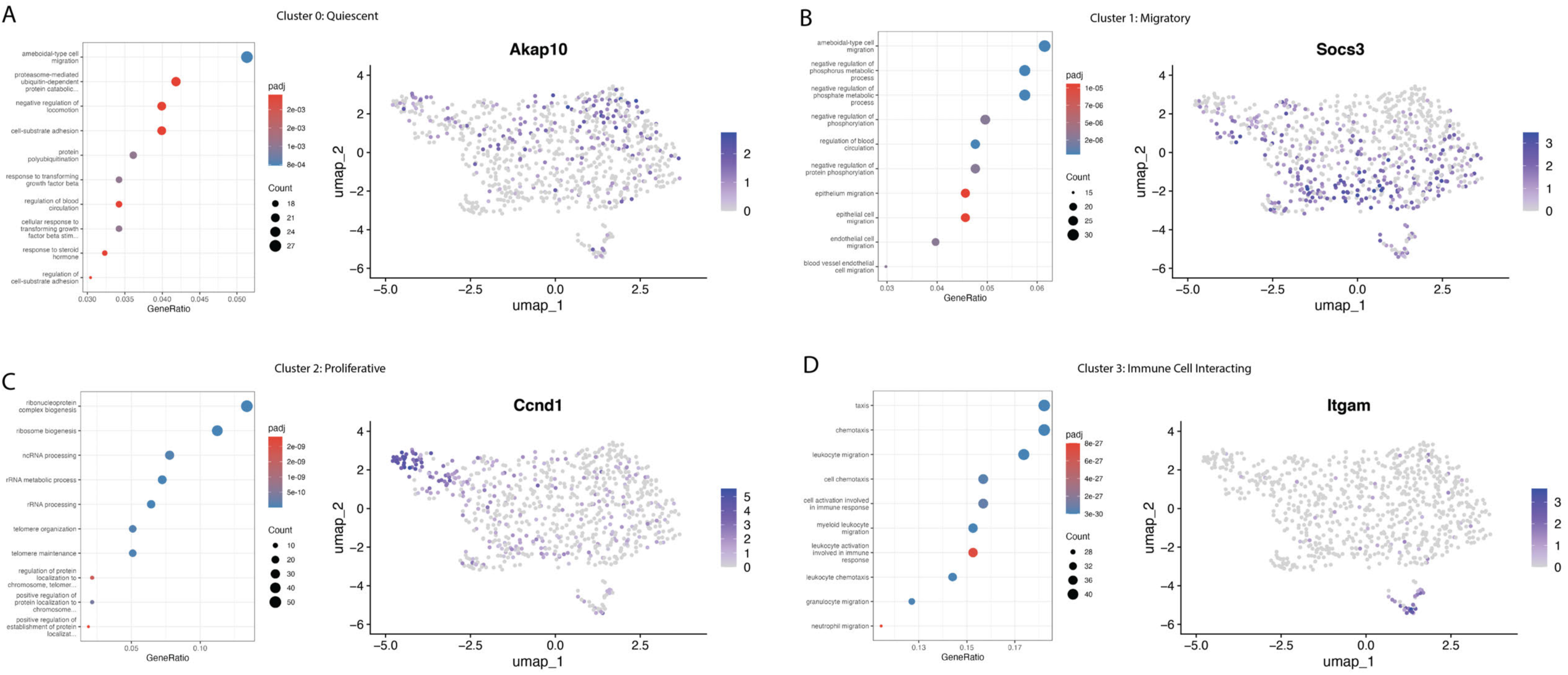
Annotation of scRNAseq EC sub-populations. Top 10 Gene Ontology (GO) biological process pathways enriched in the top 200 ranked marker genes (left panel) and feature plot of integrated endothelial cells (EC) colored by chosen cluster-specific marker gene (right panel) from cluster 0 (Quiescent; **A**), cluster 1 (Migratory; **B**), cluster 2 (Proliferative; **C**), and cluster 3 (Immune Cell Interacting; **D**).

**Supplemental Fig. 10:**
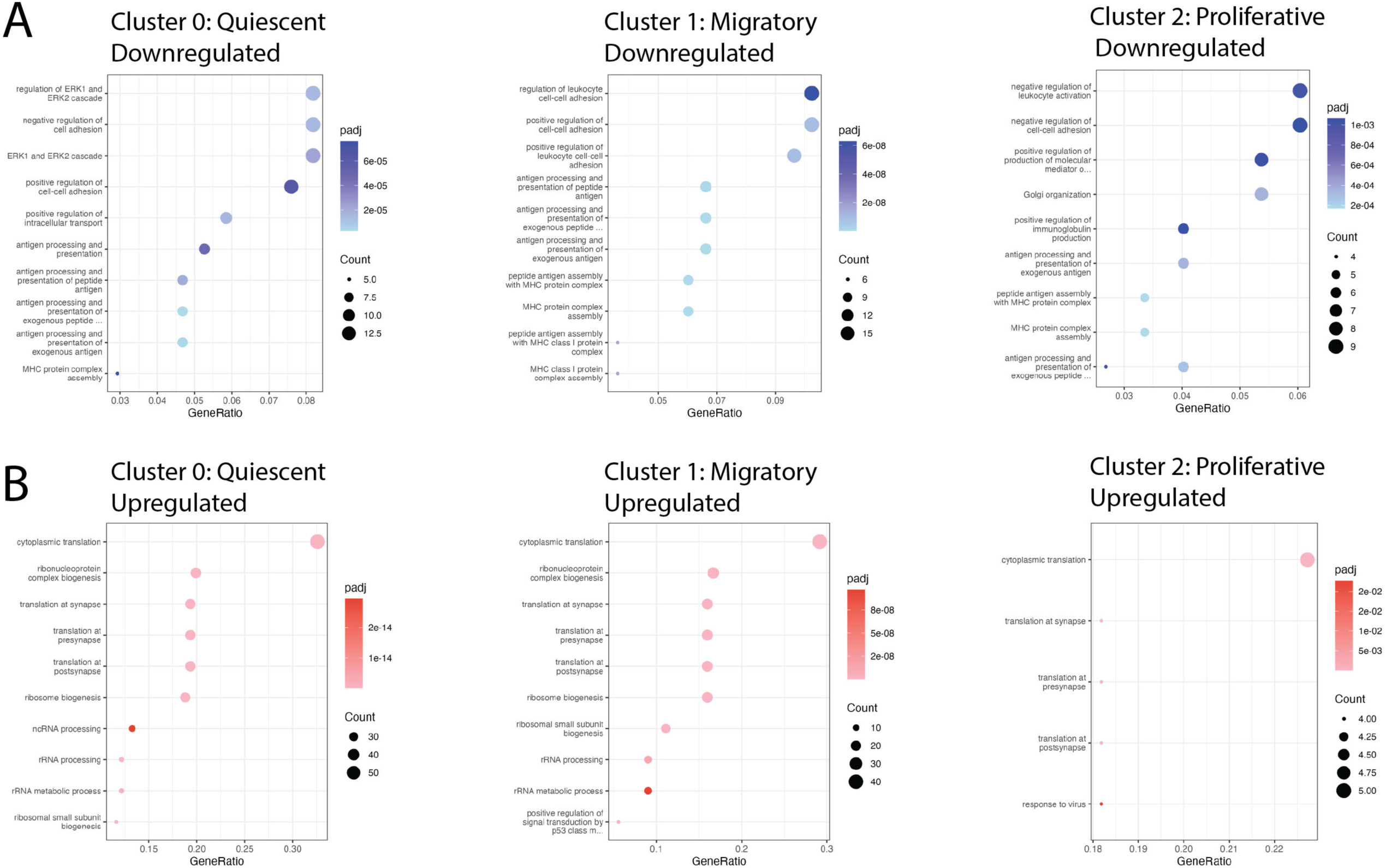
Cx43-dependent signaling pathways in injured EC. Top 10 Gene Ontology (GO) biological process pathways enriched in downregulated (Blue; **A**) and upregulated (Red; **B**) differentially expressed genes, separated by cluster, in EC-Cx43 KO cells compared to EC WT control endothelial cells.

**Supplemental Fig. 11:**
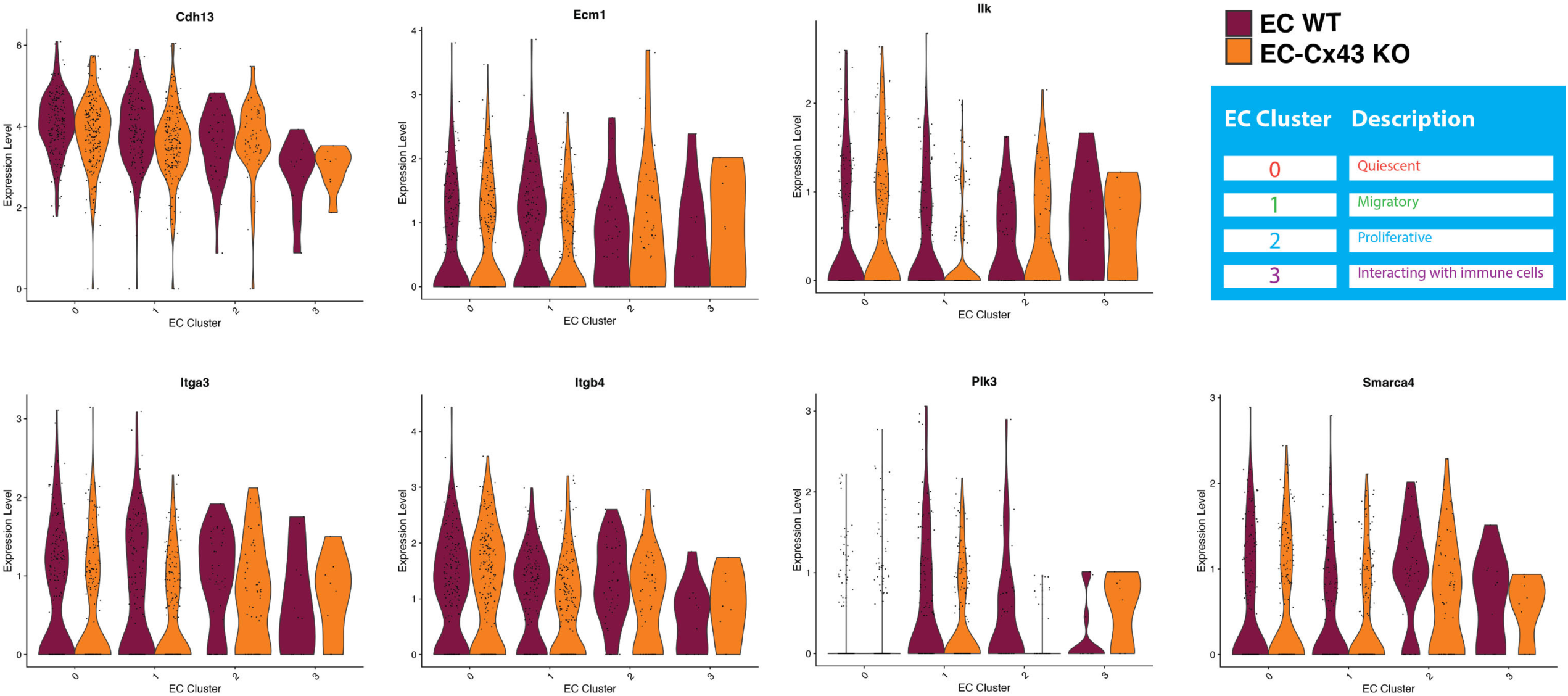
Cx43 regulates genes that control endothelial-to-mesenchymal transition (EndMT) in a cluster-specific manner. Violin plots displaying normalized gene expression levels of selected known EndMT genes in EC-WT control (maroon) and EC-Cx43 KO endothelial cells (orange) separated by cluster.

**Supplemental Fig. 12:**
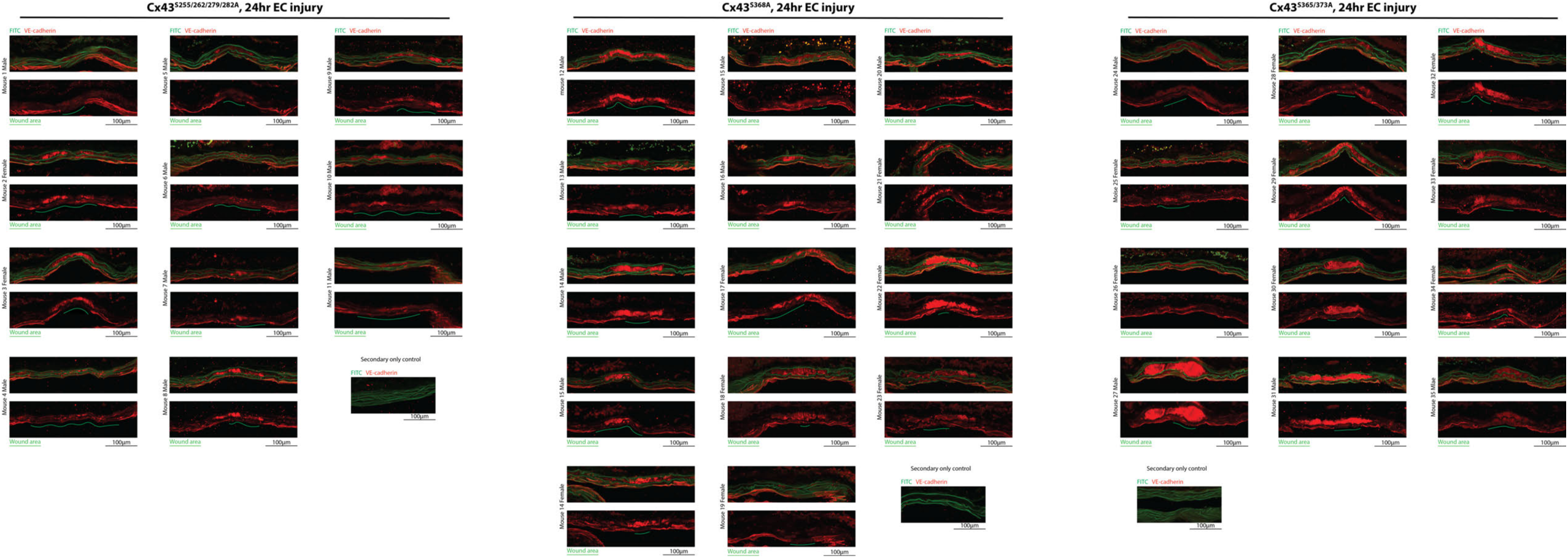
Immunofluorescent longitudinal view of Cx43^S368A^, Cx43^S255/262/279/282A^, and Cx43^S365/373A^ mouse carotids that underwent endothelial injury followed by 24 hour recovery. For each carotid, tissue autofluorescence and EC marker VE-cadherin are overlayed in the top image with VE-cadherin only in the bottom image. Green lines indicate the EC denuded area (n=9-13).

**Supplemental Table 1.**
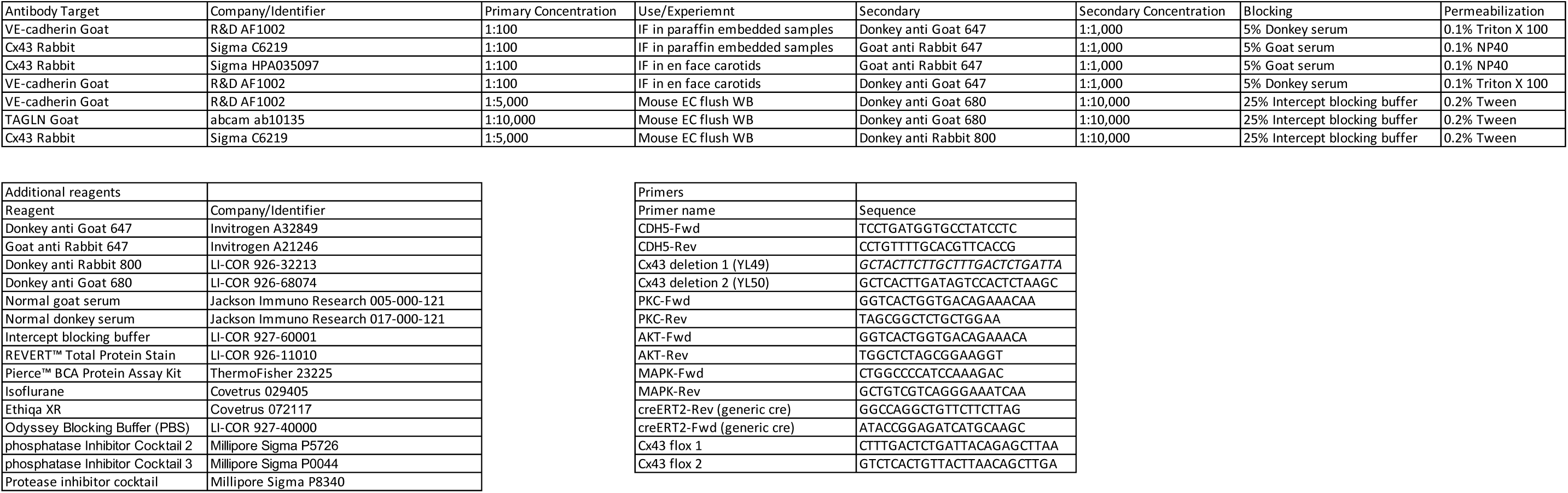

